# Bioactive coatings on 3D printed scaffolds for bone regeneration: Translation from *in vitro* to *in vivo* models and the impact of material properties and growth factor concentration

**DOI:** 10.1101/2023.10.22.560309

**Authors:** Karen. M. Marshall, Jonathan P. Wojciechowski, Vineetha Jayawarna, Abshar Hasan, Cécile Echalier, Øystein Øvrebø, Tao Yang, Janos M. Kanczler, Alvaro Mata, Manuel Salmeron-Sanchez, Molly M. Stevens, Richard O. C. Oreffo

## Abstract

Bone tissue engineering is a rapidly advancing field that seeks to develop new functional bone tissue, harnessing materials for application in bone defects which may fail to heal without intervention, as seen in critical-sized bone defects. The material properties must be developed, tailored and optimised as the environment progresses, through increasing animal size and complexity, of the target bone defect site. This study has examined the potential of a poly(caprolactone) trimethacrylate (PCL-TMA) 3D-printable scaffold with select bioactive coatings to function as a scaffold to augment bone formation. Three bioactive coatings were examined, i) elastin-like protein (ELP), ii) poly (ethyl acrylate) (PEA), fibronectin (FN) and bone morphogenetic protein-2 (BMP-2) applied sequentially (PEA/FN/BMP-2) and iii) both ELP and PEA/FN/BMP-2 coatings applied concurrently. The PCL-TMA scaffold construct was observed to be a robust scaffold material and the bioactive coatings applied were found to be biocompatible, with a significant osteogenic response from human skeletal cell populations observed *in vitro*. The PCL-TMA scaffold and bioactive coatings supported angiogenesis and displayed excellent biocompatibility following evaluation on the chorioallantoic membrane (CAM) assay. Biocompatibility was confirmed, however, no significant bone formation was detected, following examination of heterotopic bone formation in the murine subcutaneous implantation model, whereas extensive mineralisation was observed in the positive control material of collagen sponge with BMP-2. The absence of bone formation on the PCL-TMA scaffolds, *in vivo*, was potentially a consequence of the method of action of the applied coatings, the surface area of the scaffold construct for BMP-2 binding and the necessity of an appropriate *in vivo* environment to facilitate skeletal cell ingress, warranting future examination in an orthotopic bone defect model of bone tissue repair. The current studies demonstrate the development of a range of innovative scaffold constructs with *in vitro* efficacy and clearly illustrate the importance of an appropriate *in vivo* environment to validate *in vitro* functionality prior to scale up and preclinical application.

## 1. Introduction

Advances in healthcare have contributed to a global increase in population aging. However, this welcome increase in life expectancy, results in major challenges with the inability to effectively regenerate tissues, resulting in chronic illness, disability and rising healthcare expenses. The field of bone tissue engineering seeks to repair or regenerate bone tissue, harnessing innovative materials that can replace autografts (patient’s own bone), allografts (decellularized donor human bone), or a plethora of calcium phosphate-based materials currently used in clinical settings. Defects arising in bone may be due to trauma, non-union, infection, oncology treatment or congenital conditions [1]. Amputation may be required following substantial soft tissue, vascular and/or nerve damage making the limb unsalvageable [2]. Autograft remains the gold standard material for bone defect repair, as an autograft is non-immunogenic, osteoinductive (due to the presence of growth factors), osteoconductive as bone is a 3D scaffold and osteogenic given the cellular component [3-5]. The limitations to autograft application are however, donor site complications such as infection, pain and the limited volume of graft material available [6]. Thus, the ideal bone substitute material is biocompatible, bioresorbable, osteoconductive, osteoinductive, porous and with a comparable structure and strength to bone, while being easy to apply and cost effective [1].

The current work details examination of a PCL trimethacrylate (PCL-TMA) material as a biocompatible, biodegradable, 3D-printable polymer scaffold. This work builds on the research from Reznikov *et al.* where ‘octetruss’ polyamide (nylon) and titanium scaffold designs were examined, in which the more compliant nylon showed enhanced bone formation in a sheep femoral condyle defect model [7]. Bioactive surface coatings, for application in large bone defects, were examined *in vitro* and *in vivo* with potential clinical translation on the PCL-TMA octetruss scaffold. Three bioactive coatings were examined: i) elastin-like polypeptide (ELP), ii) poly (ethyl acrylate) (PEA), fibronectin (FN) and bone morphogenetic protein-2 (BMP-2) applied sequentially (PEA/FN/BMP-2) or iii) ELP and PEA/FN/BMP-2 applied concurrently on the scaffold surface.

Elastin-like polypeptides (ELPs) are artificial elastin-based polypeptides which are highly biocompatible, as ELPs are not recognised as foreign by the immune system [8]. Mineralised ELP coated scaffolds provide a novel coating based on the interaction between organic matter and inorganic crystal formation, with the formation of fluorapatite spherulites. These structures develop as prism-like microstructures and mature into macroscopic circular structures, which can encapsulate large and uneven material surfaces [9]. ELP coatings were selected given their capacity to coat implants [10] as well as their potential to mineralize while recreating the stiffness of hard tissues [11] and enhance osseointegration in bone tissue engineering [12]. Interestingly, the ELP coated PCL-TMA scaffolds displayed increased brittleness following mineralisation, limiting this combination for lower limb bone repair and therefore, the ELP coating was examined alone.

Previous work from the Salmeron-Sanchez group has demonstrated the PEA/FN/BMP-2 coating as an osteoinductive and osteoconductive material for bone regeneration [13]. PEA is non-biodegradable; however, PEA may be metabolised following application as a thin layer (<10s of nm) [13]. Fibronectin forms organised fibrils as it unfolds when in contact with PEA, enabling cell interaction with the FN adhering and aligning on the material [14]. Previous work has shown FN can bind to BMP-2, presenting BMP-2 to skeletal cells and facilitating osteogenic differentiation *in vitro* [15]. Subsequently, a mouse radial defect study illustrated enhanced bone repair with a polyimide ‘sleeve’ coated in PEA/FN/BMP-2 compared to control ‘sleeves’ [15]. The clinical use of this FN and BMP-2 growth factor interaction on PEA has been validated in veterinary cases of complicated, non-healing fractures [13, 16]. Furthermore, in comparison to other clinical reports using 0.5 mg/mL of BMP-2 solution for non-union cases, this method of binding BMP-2 to PEA/FN utilises lower concentrations of BMP-2 [17].

In the current studies, nylon scaffolds were used to test the ELP coating mineralisation efficacy due to ease of cell attachment and visibility of alizarin red staining. Extrusion printed circular PCL scaffolds with a cross-hatched porous structure were used for confirmation of the activity of the PEA/FN/BMP-2 coating on 3D scaffolds dependent on the pH of BMP-2 diluent selected (Supplementary information). The cytocompatibility of the PCL-TMA and the coatings were examined using alamarBlue™ HS cell viability reagent. The efficacy of the coatings to induce osteogenic differentiation was evaluated by ALP staining of C2C12 myoblast cells and ALP specific activity analysis as well as measuring osteogenic gene expression of human bone marrow stromal cells (HBMSCs). Subsequently, the biocompatibility, ability to integrate with vascularised tissue and to support angiogenesis was examined using the Chorioallantoic Membrane (CAM) assay. Finally, the ability of the selected bioactive scaffold constructs to induce bone formation in a heterotopic site was investigated, using the murine subcutaneous implantation model, with micro-computed tomography (µCT) quantification and histological analysis. This study illustrates the optimisation and critical analysis of *in vitro* and *in vivo* findings required on the path to clinical translation of innovative bioactive coatings.

## 2. Materials and methods

### 2.1 Materials

Reagents were purchased as follows: Ethyl acrylate (Sigma, UK), ELP with statherin sequence (SN_A_15) (Technical Proteins Nanobiotechnology, Valladolid, Spain); collagenase (Gibco, UK); human fibronectin and human recombinant BMP-2 (R&D systems, Biotechne, UK); recombinant human BMP-2 (Infuse/InductOS® Bone graft kit, Medtronic, USA); alcian blue 8X, light green SF, orange G 85% pure, Paraformaldehyde 96% extra pure, phosphomolybdic acid hydrate 80% (Acros Organics); Picrosirius Red, Van Gieson’s stain, Weigert’s Haematoxylin Parts 1 and 2 (Clintech Ltd, UK); Benzoyl peroxide, GMA solution B, JB4 solution A (Polysciences); GoTaq qPCR master mix, Herring sperm DNA, RNeasy mini prep RNA extraction kit (Promega); phosphate buffered saline (PBS), trypsin/ ethylenediaminetetraacetic acid (EDTA), Dulbecco’s Modified Eagle Medium (DMEM), Alpha Minimum Essential Medium (αMEM), penicillin-streptomycin (Scientific Laboratory Supplies, SLS); 4-Nitrophenol solution 10 nM, acetic acid, acetone, acid fushsin, alizarin red S, alkaline buffer solution, ascorbic acid-2-phosphate, beta-glycerophosphate disodium hydrate salt (βGP), cell lytic M, dexamethasone, fast violet B salts, glycine, histowax, hydrochloric acid, iodoacetamide, ipegal, L-glutamic acid, Naphthol AS-MX phosphate 0.25%, parafilm, PBS (with CaCl_2_/MgCl_2_), phenyl methyl sulphonyl fluoride, phosphatase substrate, polysorbate 80, ponceau xylidine, silver nitrate, sodium chloride, sodium hydroxide pellets, sucrose, TRIS-EDTA (TE) buffer solution (Merck, UK); Embedding capsule (TAAB Laboratories equipment); alamarBlue™ HS Cell Viability Reagent, 70 µM cell strainer, dibutyl phthalate xylene (DPX), ethidium homodimer-1, fetal calf serum (FCS), fisherbrand grade 01 cellulose general purpose filter paper, Histoclear, isopropanol, methyl benzoate, Quanti-IT™ Picogreen™ ds DNA reagent, Taqman® Reverse Transcription Kit, Vybrant™ CFDA SE Cell Tracer Kit (Thermofisher Scientific, UK); Fast green and sodium thiosulphate (VWR); Lubrithal (Dechra, UK), Isoflurane (Dechra, UK), Buprenorphine (Buprecare® multidose, Animalcare, UK) and Vetasept® sourced from MWI animal health, UK. Uncoated vacutainers, 3-way stopcock and 5/0 PDS II suture (Ethicon, USA) from NHS supply chain. All other consumables and reagents were from Sigma-Aldrich, UK.

### 2.2 Production of nylon/polyamide scaffold material

The octet-truss scaffold shape was created using Rhinoceros 3D software (McNeel, Europe, Barcelona, Spain). Nylon/polyamide scaffolds (4 mmdiameter and height, surface area minimum 90.5 mm^2^) were used as a template scaffold for analysis of ELP and mineralised coatings. Selective laser sintering (SLS) of nylon powder using an EOS FORMIGA P110 machine (Electro Optical Systems EOS Ltd., Warwick, UK) was used as previously described [7].

### 2.3 Production of PCL trimethacrylate scaffold material

PCL-trimethacrylate of this molecular weight has been synthesised and 3D printed via stereolithography (SLA) previously [18-20]. Silica gel (40-63 μm; VWR chemicals) was used as a stationary phase. ^1^H NMR and ^13^C NMR spectra were recorded on a JOEL 400 NMR spectrometer, with working frequencies of 400 MHz for ^1^H nuclei.

#### 2.3.1 Poly(caprolactone) trimethacrylate synthesis

Poly(caprolactone) triol, M_n_ = 300 Da, (50 g, 0.17 mmol, 1 eq), anhydrous dichloromethane (350 mL) and triethylamine (100 mL, 0.72 mmol, 4.3 eq) were added to a 1 L two-necked round bottom flask. The reaction was placed under a nitrogen atmosphere and then cooled in an ice-water bath for 15 minutes. A pressure-equalising dropper funnel charged with methacryloyl chloride (65 mL, 0.67 mmol, 4 eq) was attached to the round bottom flask. The methacryloyl chloride was added dropwise over approximately 3 hours. The reaction was covered with aluminium foil to protect it from light and allowed to stir and warm to room temperature (RT) overnight. The next day, methanol (50 mL) was added to quench the reaction, which was allowed to stir at RT for 30 minutes. The reaction mixture was transferred to a separating funnel and washed with 1 M aqueous hydrochloric acid solution (5x250 mL), saturated sodium bicarbonate solution (1x250 mL) and brine (1x250 mL). The organic layer was then dried with anhydrous magnesium sulphate, filtered and concentrated via rotary evaporation. The crude yellow liquid was then purified using a silica plug, with dichloromethane as the eluent. Fractions containing PCL-trimethacrylate were pooled and concentrated via rotary evaporation. The PCL-trimethacrylate was transferred to a brown glass vial and dried using a stream of air (through a plug of CaCl_2_) overnight to yield the title compound as a slightly yellow viscous liquid (82.2683 g). The PCL-trimethacrylate was supplemented with 200 ppm (w/w) of 4-methoxyphenol (MEHQ) as an inhibitor (16.34 mg).

^1^H NMR (400 MHz, CDCl_3_) δ 6.14 – 6.04 (m, 3H), 5.63 – 5.50 (m, 3H), 4.18 – 4.00 (m, 9H), 2.36-2.32 (m, 3H), 1.94 (m, 9H), 1.75 – 1.47 (m, 9H), 1.47 – 1.32 (m, 2H), 1.02 – 0.83 (m, 3H).

The characterisation data agrees well with that previously reported, however, with an improved degree of functionalisation (>95%).

#### 2.3.2 3D printing of PCL-trimethylacrylate

PCL-trimethacrylate octetruss scaffolds (5 mm diameter and height, surface area 143.4 mm^2^), denoted PCL-TMA scaffolds, were printed using masked (SLA) 3D printing on a Prusa SL1. The resin was prepared for 3D printing by first dissolving 0.1% (w/w) 2,5-thiophenediylbis(5-tert-butyl-1,3-benzoxazole) (OB+) as a photoabsorber in the PCL-TMA by stirring at RT for 1 hour. Finally, 1.0% (w/w) diphenyl(2,4,6-trimethylbenzoyl)phosphine oxide (TPO-L) as a photoinitiator was added to the resin.

After printing, the scaffolds were rinsed with ethanol and removed from the build plate. Scaffolds were sonicated in ethanol (5x5 minutes) and allowed to dry for 15 minutes at RT. The scaffolds were post-cured using a Formlabs Form Cure for 60 minutes at RT. After post-curing, the scaffolds were soaked into ethanol overnight at RT on a rocker (100 rpm), rinsed with ethanol (3x) and allowed to dry at RT before being ELP and/or PEA coated and EO sterilised, or left uncoated and EO sterilised.

### 2.4 HBMSC isolation and culture

#### 2.4.1 Isolation and culture of HBMSCs

Human bone marrow samples were collected from patients undergoing hip replacement surgery, identified only by sex (male (M) or Female (F)) and age (e.g., F60) to maintain confidentiality, with approval of the University of Southampton’s Ethics and Research Governance Office and the North West-Greater Manchester East Research Ethics Committee (18/NW/0231). In a class II hood, under sterile conditions, 5-10 mL alpha-Minimum Essential Medium (α-MEM, Lonza, UK) or Dulbecco’s Modified Eagle Medium (DMEM, Lonza, UK) was added to the universal tube of marrow and shaken vigorously to extract the HBMSCs. A 3 mL sterile Pasteur pipette was used to remove the supernatant media/cellular debris mix to a 50 mL falcon tube and washing was repeated until the bone was light pink/white in colour. The cell suspension was centrifuged (272 g Heraeus mega 1.0R centrifuge) for 5 minutes. The supernatant was removed, the pellet resuspended in α-MEM or DMEM and passed through a 70 µM cell strainer (Fisher Scientific, UK) to remove bone and fat debris. The suspension was centrifuged and the supernatant poured off. The pellet was resuspended in basal media (α-MEM, 10% fetal calf serum (FCS), 1% penicillin-streptomycin (P/S)) and the HBMSCs were cultured in T175 flasks at 37 °C in 5% CO_2_/balanced air until approximately 80% confluent. Collagenase (2% solution and or 0.22 IU/mg) was used prior to trypsin solution (1x concentration (Stock Trypsin/EDTA (10X), includes 1,700,000 U\L trypsin 1:250 and 2 g/L Versene® (EDTA)), for passaging and seeding onto scaffolds.

Osteogenic media consisted of α-MEM, 10% FCS, 1% P/S, ascorbate-2-phosphate 50 mM (2 µL/mL), dexamethasone 10 μM (1 µL/mL). Mineralisation media consisted of α-MEM, 10% FCS, 1% P/S, ascorbate-2-phosphate 50 mM stock (1 µL/mL), dexamethasone 10 μM stock (1 µL/mL) and 2 μM beta-glycerophosphate (βGP) 1 M stock (2 μL/mL). Mineralisation media was used for alizarin red staining experiments only. All media was changed every 3-4 days.

#### 2.4.2 Cell seeding onto scaffolds

2.5 ×10^4^ Passage 2 (P2) HBMSCs per scaffold were used for cell viability/alamarBlue™ HS experiments and 5×10^4^ Passage 1 (P1) HBMSCs were used for biochemistry and molecular experiments (**Supplementary Figure 1**). Each PCL-TMA octetruss scaffold was added individually to a 2 mL Eppendorf tube and 500 µL cell suspension (α-MEM, 1% P/S, without FCS) was added. The Eppendorf tubes were placed in a 50 mL falcon tube (6 per tube) for positioning horizontally on the MACSmix™ Tube Rotator enabling a maximum of 24 scaffolds to be seeded per rotator machine. The Eppendorf tubes were fully sealed and therefore the air available to cells was that only within the tubes. The media maintained the normal pink colour throughout cell seeding, indicating the pH of the media was unaltered. FCS was added at approximately 10% (55 µL) to each tube after 3 hours. After 24 hours, each scaffold was moved to individual wells of a 24 well plate containing 1.5 mL basal or osteogenic media. Culture was performed at 37 °C in 5% CO_2_/balanced air, with media changed every 3 days. Each scaffold coating type/media condition was set up in triplicate.

#### 2.4.3 Culture of HBMSCs on nylon scaffolds for alizarin red staining assay

The uncoated scaffolds were placed in 50 mL falcon tubes and submerged in 70% ethanol for sterilisation. The tube was centrifuged at 1700 g in the Heraeus mega 1.0R centrifuge for 3 minutes at RT. The ethanol was removed using a 3 mL Pasteur pipette and replaced with phosphate buffered saline (PBS 1x). Centrifugation was repeated with fresh PBS, followed by replacement with α-MEM for rinsing twice again. The ELP coated and ELP coated scaffolds which had been mineralised were placed in 10 mL falcon tubes and submerged in 70% ethanol. The tubes were placed on a rotator at 180 rpm for 30 minutes. The ethanol was removed and replaced with PBS twice, followed by α-MEM twice, each for 15-30 minutes on the rotator at RT. 5×10^4^ P1 HBMSCs were seeded onto the scaffolds using the MACSmix™ rotator as described. After 24 hours, the scaffolds were moved to a 24 well plate and 1.5 mL basal or mineralisation media was added. The culture was continued for 27 days until staining with alizarin red S stain. Control scaffolds with no cells seeded were established to determine staining due to exogenously applied mineral during coating of the scaffold, compared to mineralised matrix produced by the HBMSCs.

### 2.5 Coating of the scaffold material

#### 2.5.1 PEA/FN/BMP-2 coating of the scaffold material

PEA coating of materials was performed as previously described [13]. Briefly, the scaffolds were treated in air plasma for 5 minutes before being exposed to monomer plasma. Plasma polymerization was carried out in a custom-made capacitively coupled plasma installation for low-pressure plasma in a 15-L T-shaped reactor made of borosilicate glass and stainless-steel end plates sealed with Viton O-rings. A vacuum was produced by a rotary pump or a scroll pump (both BOC Edwards, UK), with working pressures for the monomer plasma from 0.15 to 0.25 mbar. The plasma was initiated via two capacitively coupled copper band ring electrodes located outside of the reactor chamber and connected to a radiofrequency generator (Coaxial Power System Ltd.). The monomer pressure was controlled via speedivalves (BOC Edwards, UK) and monitored with a Pirani gauge (Kurt J. Lesker). PEA was applied on the material surfaces for 15 minutes at a total radiofrequency power of 50 W. The samples were sterilized afterwards using ethylene oxide (EO).

The application of the BMP-2 coating was optimised by investigating different diluents and BMP-2 manufacturers to ensure the biological activity of the BMP-2 protein (**Supplementary Table 1, Supplementary** Figures 2, 3 and 4). For experiments with the PEA/FN/BMP-2, or ELP/PEA/FN/BMP-2 coated PCL-TMA scaffolds, FN and InductOs® BMP-2 were diluted in PBS with calcium chloride (CaCl_2_) and magnesium chloride (MgCl_2_) added. A maximum of three scaffolds were placed into a non-coated 11 mL vacutainer and 1 mL of FN solution (20 µg/mL) was added. A vacuum was created and after 1 hour in the sterile hood at RT, the FN solution was removed. PBS (containing CaCl_2_/MgCl_2_) was added at 1-2 mL per vacutainer to rinse off non-bound FN and repeated once. The scaffolds were handled with sterile forceps and transferred to new vacutainers. 1 mL of BMP-2 solution (100 ng/mL for *in vitro* experiments or 5 µg/mL for CAM assay and subcutaneous implantation experiments) was added to each tube. The formation of a vacuum was repeated. After 1 hour at RT, the BMP-2 solution was removed and the scaffolds rinsed twice in PBS (containing CaCl_2_/MgCl_2_) prior to use.

#### 2.5.2 ELP and mineralised coating of the scaffold material

ELP coating of scaffolds was based on previous published methods [11]. In brief, lyophilized ELP powder was dissolved in solvent mixture of DMF/DMSO (at 9/1 ratio) to prepare 5% (w/v) ELP solution followed by addition of hexamethyl diisocyanate (HDI) crosslinker (cross-linker to lysine ratios of 12/1). 3D printed nylon or PCL-TMA scaffolds were immersed in the ELP solution for 10–15 seconds and left to dry overnight at RT (22 °C) inside a glovebox (BELLE Technology, UK) maintained at a humidity <20%. Dry ELP coated scaffolds were washed several times with deionised water (dH_2_O) to remove excess HDI and were termed as ELP coated scaffolds [12].

Mineralizing solution was prepared using reported methodology [11]. Briefly, hydroxyapatite powder (2 mM) and sodium fluoride (2 mM) were dissolved in dH_2_O by dropwise addition of nitric acid (69%, v/v) into the solution until it became clear. The pH of the solution was adjusted to 6.0 using 30% (v/v) ammonium hydroxide solution. To create ‘mineralised’ scaffolds, ELP coated scaffolds were incubated in the mineralizing solution (pH 6) at 37 °C for 16 days. Post mineralisation, scaffolds were washed several times with dH_2_O, air dried and stored at RT until use [12]. The ELP coating and mineralisation were confirmed on the PCL-TMA scaffold surface using scanning electron microscopy [12].

For the ELP/PEA/FN/BMP-2 coated scaffolds, the ELP coating was applied, followed by PEA and subsequent EO sterilisation. Finally, the FN and BMP-2 were adhered to the scaffolds as described in section 2.5.1.

### 2.6 Assessment of cytocompatibility of PCL-TMA and coatings

#### 2.6.1 alamarBlue™ HS Cell Viability assay

For cell viability experiments alamarBlue™ HS Cell Viability Reagent was added to basal media at 10% (v/v) concentration. Fluorescence measurements were taken at day 1 (when the PCL-TMA scaffolds were removed from the Eppendorf tube after 24 hours of cell seeding) and on day 14 (each PCL-TMA scaffold was moved to a new 24 well plate to ensure only the cells adhered to the scaffold were quantified). A 1 mL aliquot of media/alamarBlue™ HS mix was added to each well containing a scaffold and to 3 wells with no scaffold as background measurements and incubated for 4 hours at 37 °C in 5% CO_2_/balanced air. After 4 hours, 100 µL samples were taken from each well and plated in triplicate in a black 96 well plate. The fluorescence was measured using the GloMax® Discover Microplate Reader (Promega, UK) at Green 520 nm excitation and 580-640 nm emission and the average background measurement was subtracted from each sample well.

#### 2.6.2 Labelling of live and dead HBMSCs

Vybrant™ CFDA SE Cell Tracer 10 μM and 5 µg/mL Ethidium homodimer-1 in PBS, were used to label live and dead cells respectively on PCL-TMA scaffolds at day 1 and day 14 post alamarBlue™ HS analysis. Media/alamarBlue™ HS was removed and scaffolds washed twice in PBS. A 1 mL aliquot of labelling solution was added to cover the scaffolds/cells and incubated for 15 minutes at 37 °C in 5% CO_2_/balanced air. The labelling solution was removed and replaced with α-MEM/1% P/S/10% FCS and culture continued for 30 minutes. The samples were imaged under fluorescence microscopy using the FITC filter (excitation 485 nm, emission 515 nm) for live cells and RHODA/TRITC filter (excitation 510-560 nm, emission 590 nm) for dead cells, with a Zeiss Axiovert 200 microscope and Axiovision 4.2 imaging software.

### 2.7 Biochemistry assays for HBMSC differentiation analysis

#### 2.7.1 Alkaline Phosphatase Specific Activity measurement

Placement of scaffolds in basal or osteogenic media after 24 hours of cell seeding was determined as day 0. Therefore, day 1 was after 24 hours of culture in basal or osteogenic media and so on until the day 7 when the HBMSCs were lysed. HBMSCs attached to PCL-TMA scaffolds were washed twice in PBS and the scaffolds were transferred to individual 2 mL Eppendorf tubes. A 200 µL aliquot of Cell lytic M was added to cover the scaffold and left in contact for 15 minutes at RT with vortexing performed three times every 5 minutes. The Eppendorf tubes were stored at -80 °C.

ALP activity was measured using a colourimetric absorption assay. P-nitrophenol phosphate (pNPP) production was measured against standards. A 10 µL aliquot of cell lysate was transferred to a clear 96 well plate and 90 µL of substrate was added. The plate was incubated at 37 °C until a yellow colour change was noted and the time recorded for this change. A 100 µL aliquot of 1M sodium hydroxide solution was added to stop the reaction, prior to reading absorbance on the GloMax® Discover (spectrophotometer) at 405 nm. The same centrifuged cell lysate samples were used as for ALP quantification. PicoGreen™ was diluted 1/200 in TE (1x) buffer and added to all wells, including standards. The plate was left on the benchtop at RT for 5 minutes prior to the quantity of DNA being measured using a fluorescence assay at blue 475 nm excitation and 500-550 nm emission, in a GloMax® Discover Microplate Reader (Promega, UK). Results expressed as nanogram (ng) of DNA. ALP specific activity was calculated as ALP produced per ng of DNA as nmol pNPP/ng DNA/hr.

#### 2.7.2 Alizarin Red staining of nylon scaffolds for mineralisation

All scaffolds were washed with PBS twice and fixed with 1 mL 4% paraformaldehyde (PFA) for 15 minutes at RT and rinsed 3 times with dH_2_0. Alizarin red S solution (40 mM, 1 mL) was added to each well for 1 hour at RT on the horizontal shaker at 180 rpm. The solution was discarded and the scaffolds rinsed in dH_2_0 until the stain was no longer released. Images were taken using a Canon G10 camera attached to a stemi 2000-c stereomicroscope (Zeiss, UK) with identical magnification and camera brightness settings to allow subjective comparison of staining intensity.

### 2.8 Osteogenic gene expression analysis

#### 2.8.1.1 RNA extraction

RNA extraction was performed using the Promega ReliaPrep™ RNA Cell Miniprep System instructions. Scaffolds were washed in PBS and transferred to 2 mL Eppendorf tubes with BL-TG buffer (250 µL) for 15 minutes to lyse the cells prior to storage at -80 °C. The tubes were thawed and isopropanol (85 µL) added, prior to brief vortexing. The lysate was transferred to a minicolumn within a collection tube. The column was washed in a series of steps as per the kit instructions and the final RNA eluted with nanopure water (15 µL). RNA quantification and purity measurement were performed using the Nanodrop 2000 spectrophotometer.

#### 2.8.1.2 Reverse transcription

RNA quantity for conversion to cDNA was standardised for all samples with dilution in nanopure water to a volume of 9.6 µL in a 0.2 mL PCR reaction tube. The reagents of the TaqMan® Reverse Transcription (RT) Kit were mixed in quantities instructed to make RT master mix and 10.4 µL was added to water/RNA mix to give a final 20 µL reaction volume. The tubes were placed in a SimpliAmp thermal cycler (Applied Biosystems, UK) set to 10 minutes at 25 °C for primer incubation, 30 minutes at 37 °C for reverse transcription and 5 minutes at 95 °C for the reverse transcription inactivation and 4 °C to hold samples until retrieval for storage at -20 °C.

#### 2.8.1.3 Quantitative Polymerase Chain Reaction (qPCR)

Quantitative qPCR was performed using GoTaq PCR master mix. The master solution was made from 10 µL GoTaq PCR master mix, 0.75 µL of forward primer, 0.75 µL of reverse primer and 7.5 µL ultrapure water. The final 20 µL reaction volume was made of 2 µL of cDNA sample and 18 µL of master solution in a 96 well PCR plate, sealed and centrifuged briefly before analysis using 7500 Real Time PCR system (Applied Biosystems, UK). The resulting data were collected using the AB7500 SDS Software (Applied Biosystems, UK). Ct values for each sample were normalised to the housekeeping gene β-actin. Relative-expression levels of each target gene were calculated using the ΔΔ C_t_ method (cycle threshold) method. The uncoated scaffold cultured in basal media was used as an internal relative reference for the other scaffolds and media types. Each sample was a biological triplicate and plated out once each.

### 2.9 Statistical analysis

alamarBlue™ and biochemistry results were established with biological triplicates, with triplicate readings taken from each sample and the average of the triplicate readings used for statistical analysis. Molecular experiments were run using biological triplicates with one reading taken from each sample. Statistical analysis was performed comparing uncoated scaffolds to each coated type and in basal and osteogenic conditions where applicable. Statistical analysis of the molecular results used the ΔCT values, while the graphs displayed show the 2^-ΔΔCT^ values to display fold change in relation to the uncoated scaffolds cultured in basal media. Analysis and graphical presentation were performed using GraphPad Prism 9, version 9.2.0. P values <0.05 were considered significant. Graphical representation of significance as follows: ns is no significant difference, *p<0.05, **p<0.01, ***p<0.001, ****p<0.0001. All data presented as mean and standard deviation (S.D.).

### 2.10 Chorioallantoic Membrane assay

Descriptions are extrapolated from Marshall *et al.* [21]. All procedures were performed with prior received ethical approval and in accordance with the guidelines and regulations stated in the Animals (Scientific Procedures) Act 1986, however a UK Personal Project License (PPL) was not required as the eggs were used only until embryonic day (ED) 14 when the experiment ended. Briefly, fertilised hens (*Gallus gallus domesticus*) eggs (Medeggs, Henry Stewart & Co., Lincolnshire) were incubated in a humidified (60%), warm (37 °C) incubator (Hatchmaster incubator, Brinsea, UK) at embryonic day (ED) 0. The eggs were incubated in a horizontal position, on a rotating pattern (1 hour scheduled rotation), prior to albumin removal at ED 3 (**Supplementary Figure 5**). The minimum number of eggs required to determine a significant difference (p<0.05) between groups was calculated with power 80%. Typically, an n=6 for each condition (to allow predominantly for non-developing eggs) was used.

#### 2.10.1 Scaffold preparation and implantation of materials

All PCL-TMA scaffolds (uncoated, ELP coated and/or PEA coated) were EO sterilised. The PEA scaffolds and ELP/PEA coated scaffolds were coated with FN (20 μg/mL) and BMP-2 (5 μg/mL) solutions immediately prior to use. Scaffolds were implanted at ED 7. Descriptions are extrapolated from Marshall *et al* [21]. In brief, eggs were candled to check viability and a No. 10 scalpel blade was used to create a 0.5 cm by 0.5 cm window. The white inner shell membrane was peeled away and the materials placed onto the CAM. Parafilm was used to cover the window and labelled tape applied, parallel to sides of the egg, to hold the parafilm in place. The eggs were placed horizontally within an egg incubator at 37 °C and 60% humidity without rotation.

#### 2.10.2 Analysis of results

Samples were harvested at ED 14 of incubation with methodology extrapolated from Marshall *et al* [21]. Blinding of the assessor was performed. The window was opened digitally and with forceps to image the scaffold/CAM using a stemi 2000-c stereomicroscope (Zeiss, UK), for illustrative, recording purposes only. Quantification of angiogenesis was performed using the Chalkley eyepiece graticule scoring method. Three separate counts were made and the average score was calculated for each egg. Biocompatibility was assessed by counting live, viable and developed chicks and any dead/deformed chicks. Thereafter, the scaffold and 1 cm diameter of surrounding CAM tissue was collected and placed into 2 mL 4% PFA in a 24 well plate for 72 hours at 4 °C followed by exchange of PFA for 70% ethanol. Processing of the scaffold/tissue, glycidyl methacrylate (GMA) resin embedding and subsequent histology followed. The chick was euthanised by an approved schedule 1 method at ED 14. Data was analysed using GraphPad Prism 9, software version 9.2.0. Data was presented as the mean and standard deviation (S.D.). P values <0.05 were deemed significant. Set up used n=6 eggs per condition, but only the mean and S.D. of the Chalkley score results from formed/viable eggs or those able to be counted were included.

### 2.11 Murine subcutaneous implantation study

#### 2.11.1 Mouse type and housing

All procedures were performed in accordance with institutional guidelines and with ethical approval and under PPL P96B16FBD, in accordance with the regulations in the Animals (Scientific Procedures) Act 1986 and using the ARRIVE guidelines. Nine, adult, male MF-1 wild type mice, bred on site and group housed in individually ventilated cages (IVCs) were used. Mice had access to *ad libitum* standard pellets and water.

#### 2.11.2 PCL-TMA scaffold and collagen sponge with BMP-2 preparation for subcutaneous implantation

The PCL-TMA scaffolds were uncoated, ELP and/or PEA coated followed by EO sterilisation. The PEA coated and ELP/PEA coated scaffolds had FN and BMP-2 applied immediately prior to use. Three scaffolds were used per vacutainer therefore theoretically 6.6 µg FN and 1.6 µg BMP-2 could be adhered to each scaffold if coated equally. The positive control was a collagen sponge disc (4 mm diameter, 2 mm high) soaked with 5 µg of BMP-2 within 15 µL of InductOs® buffer solution (333.33 µg/mL) (**Supplementary Table 1**) immediately prior to use. An n=6 mice per coated scaffold group and n=9 mice in the uncoated PCL-TMA scaffold and collagen sponge/BMP-2 control groups were used.

#### 2.11.3 Subcutaneous implantation surgical procedure

General anaesthesia was induced with volatile Isoflurane (5%) and 100% oxygen (0.8 L/min flow rate) and this was continued for the maintenance of a surgical plane of anaesthesia at approximately 1.5-2.5% isoflurane. Once anaesthetised, Lubrithal eye gel was applied, ear marking performed for identification, the dorsum of the mouse was shaved and chlorhexidine/alcohol solution (Vetasept® Clear Spray) applied to the dorsum and allowed to dry. Buprenorphine (Buprecare® multidose, 0.05 mg/kg) was administered subcutaneously. Four dorsal 5 mm incisions were made (2 on each side over the shoulder and flank regions) and the skin elevated using blunt dissection, with a scaffold or collagen sponge/BMP-2 implanted into each subcutaneous pouch. The skin incisions were closed in a simple interrupted, horizontal mattress suture pattern with absorbable 5/0 PDS (Ethicon, USA), with the knots towards midline. The mice were recovered inhaling 100% oxygen, whilst being kept warm with a heat-pad underneath the surgical drape. After observed movement, the mouse was transferred to a heating box (Thermacage™, datesand group, UK) prewarmed to 30 °C until normal mouse activity resumed (eating, walking, grooming) at which point the mouse was returned to IVC housing.

#### 2.11.4 Micro-CT procedure and analysis of results

Micro-CT (µCT) was performed using a MILabs OI-CTUHXR preclinical imaging scanner (Utrecht, The Netherlands). The scaffolds/collagen sponges were scanned *in vivo* at weeks 2, 4, 6 and the excised materials only at week 8. General anaesthesia was induced using 5% isoflurane in an induction box, the mouse moved to the imaging bed and maintained at 1.5-2.5% isoflurane throughout imaging, with the oxygen flow rate constant at 0.8-1.0 L/minute. BioVet software connected to the Milabs scanner, permitted monitoring of respiratory activity (minimum 60 breaths per minute) and setting of the temperature of the scanning bed to 34 °C. Lubrithal eye gel was applied before imaging to prevent eye desiccation throughout the scan. Mice were imaged using 3 bed positions to µCT scan from the neck to the base of the tail, with a total scan time of approximately 15 minutes. Mice were recovered on the warm imaging bed followed by the heating box. After 8 weeks, to obtain higher resolution images; scaffolds were retrieved from the mice post-mortem and scanned in a specialised sample holder. A density phantom, as a reference for quantification of bone density, was scanned using the same parameters (**Supplementary Table 2**). μCT reconstructions were obtained via MILabs software (MILabs-Recon v. 11.00). Formation of bone was assessed using Imalytics Preclinical software v3.0 (Gremse-IT GmbH). A gauss filter of 0.5 and bone density threshold for analysis was set equal to the average result from the lower density bone phantom. Data from replicates were analysed using GraphPad Prism 9, version 9.2.0. P values <0.05 were considered significant with all data presented as mean and standard deviation (S.D.).

### 2.12 Histology

#### 2.12.1 GMA resin embedding and sectioning of samples

Tissue samples with scaffolds/collagen sponge were fixed in 4% PFA at 4 °C for up to 7 days and stored in 70% ethanol at 4 °C. The PCL-TMA scaffolds and collagen sponge/BMP-2 samples were placed in glass vials of ice-cold acetone containing 2 mM phenyl methyl sulphonyl fluoride and 20 mM iodoacetamide overnight at -20 °C. The solution was removed and acetone added for 15 minutes at RT, this was removed and methyl benzoate added for 15 minutes at RT. The methyl benzoate was removed and a solution of 5% methyl benzoate in glycerol methacrylate (GMA solution A) was added and incubated at 4 °C for 6 hours total, with fresh solution added after 2 and 4 hours. After 6 hours, this solution was removed and embedding solution was made by adding 10 mL GMA solution A to 70 mg benzoyl peroxide and swirling until the powder dissolved. A 250 µL aliquot of GMA solution B was added and swirled by hand until the colour became off-yellow. The samples were placed into labelled plastic embedding tubes and the resin pipetted to completely fill each tube, with an ellipse at the top and the lid closed firmly. The samples were placed in the fridge at 4 °C with a weight on top to keep the lids tight to exclude any air and left for at least 48 hours. The samples were transferred to a plastic box with silicone granules and stored at -20 °C. Samples were sectioned at 10 µm using a Reichert-Jung microtome with tungsten carbide Micron blade and floated in water onto glass slides. The slides were dried at RT until staining.

#### 2.12.2 Histology staining

Staining was performed without prior rehydration of the sample sections and was performed with Alcian Blue and Sirius Red, Goldner’s Trichrome, Alizarin Red and Von Kossa stains followed by blotting to dry and mounting with dibutyl phthalate xylene (DPX) and a glass coverslip and allowed to dry. Histological samples were imaged using the Zeiss Axiovert 200 digital imaging system using bright field microscopy with the halogen bulb. Images were captured using the Axiovision 4.2 imaging software.

#### 2.12.2.1 Alcian blue/Sirius red staining

Weigert’s Haematoxylin was applied for 10 minutes to stain the cell nuclei. Excess stain was removed by rinsing 3 times in acid/alcohol (5% HCl/70% ethanol) followed by dripping on water for 5 minutes. 0.5% Alcian blue 8GX in 1% acetic acid was applied for 10 minutes. Slides were covered in 1% molybdophosphoric acid for 10 minutes, followed by rinsing in water prior to staining with 1% Picrosirius Red (Sirius Red) for 1 hour and excess stain was rinsed off with water.

#### 2.12.2.2 Goldner’s Trichrome staining

The method followed was identical to Alcian blue/Sirius Red staining to the point of acid/alcohol use and rinsing with water for 5 minutes. Ponceau Acid Fuchsin/Azophloxin was applied for 5 minutes followed by a 15 second wash with 1% acetic acid dripped onto the slide. Phosphomolybdic acid/Orange G was applied for 20 minutes followed by another 15 second wash with 1% acetic acid. Light Green was applied for 5 minutes followed by the third 15 second wash with 1% acetic acid.

#### 2.12.2.3 Von Kossa staining

The slides were incubated with 1% silver nitrate under UV light for 20 minutes. Slides were washed using water and incubated with 2.5% sodium thiosulfate for 8 minutes, washed with water and counterstained with Alcian blue for 1 minute. The slides were washed in water and van Gieson’s stain was applied for 5 minutes.

#### 2.12.2.4 Alizarin red/Light green staining

Alizarin red S (40 mM) stain solution was applied for 2 minutes. The slide was blotted on paper towel and light green stain applied for 2 minutes as a counterstain. This was blotted off and acetone followed by acetone/Histoclear (50:50, (v/v)) followed by Histoclear were applied for 30 seconds each and the excess allowed to evaporate prior to mounting.

## 3. Results

### 3.1 In vitro cell viability and osteogenic differentiation on PCL-TMA scaffolds

#### 3.1.1 The PCL-TMA material and bioactive coatings were cytocompatible with HBMSC attachment and growth over 14 days

Cell number was measured using alamarBlue™ HS reduction as a surrogate marker (**Supplementary Figure 6**). Preliminary studies confirmed the PEA/FN/BMP-2 coating was cytocompatible on PCL-TMA scaffolds and the PEA/FN/BMP-2 bioactive coating was observed to increase cell adhesion, as determined by enhanced fluorescence values obtained at day 1, although this did not reach significance (**Supplementary Figure 7**). The ELP, PEA/FN/BMP-2 and ELP/PEA/FN/BMP-2 coatings were observed to be cytocompatible with an increase in alamarBlue™ HS fluorescence results (**Figure 1 A**) and cell number at day 14, as observed by fluorescent staining of the cells (**Figure 1 B**). The PEA/FN/BMP-2 coating on the PCL-TMA scaffold, or in conjunction with the ELP coating, were observed to enhance cell adhesion to the PCL-TMA scaffold material after 24 hours as evidenced by increased alamarBlue™ HS fluorescence.

**Figure 1:**
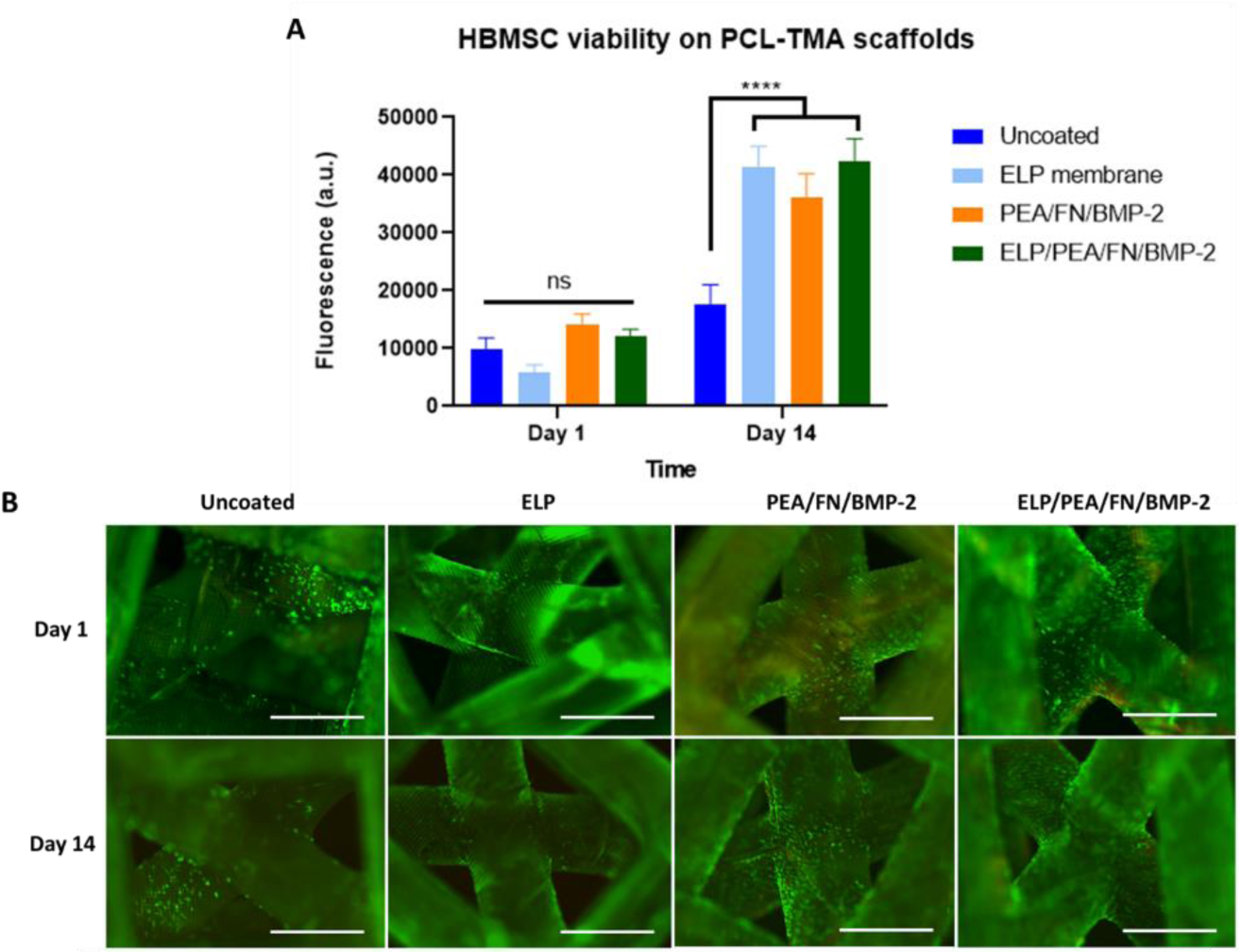
HBMSC viability and live (green)/dead (red) cell labelling on uncoated, ELP, PEA/FN/BMP-2 and ELP/PEA/FN/BMP-2 coated PCL-TMA scaffolds. (A) alamarBlue™ HS fluorescence results of ELP, PEA/FN/BMP-2 and ELP/PEA/FN/BMP-2 coated PCL-TMA scaffolds show no significant difference between uncoated PCL-TMA and coated scaffolds, however all coating variations showed a significantly increased fluorescence result at day 14 compared to uncoated PCL-TMA. 2-way ANOVA with Dunnett’s multiple comparisons test, n=3, mean and S.D. shown, ns; non-significant, ****p<0.001. (B) The cell number and location of cells adhered to the scaffold initially at day 1 and subsequent increase in cell coverage at day 14, as seen by fluorescent labelling of cells. Scale bar 1 mm.

#### 3.1.2 Osteogenic differentiation of HBMSCs in response to ELP coated, PEA/FN/BMP-2 and ELP/PEA/FN/BMP-2 coatings on 3D scaffolds

The ELP coated, PEA/FN/BMP-2 coated, or combined coating did not significantly increase ALP activity of HBMSCs in basal or osteogenic culture conditions compared to HBMSCs cultured on uncoated PCL-TMA scaffolds (**Figure 2 A**). Due to these unexpected ALP specific activity results following HBMSC culture with PEA/FN/BMP-2 coated PCL-TMA scaffolds, an increased concentration of BMP-2 was investigated. Preliminary scaffold material, nylon, was used as the base for the PEA/FN/BMP-2 coating application rather than PCL-TMA, due to ease of printing and use of the nylon material in optimisation experiments. Nylon PEA/FN coated scaffolds were examined with 100 ng/mL and 5 μg/mL BMP-2 and compared to the uncoated scaffold, given a 5 μg/mL BMP-2 concentration was typically used for the *in vivo* CAM assay and murine subcutaneous studies. Addition of BMP-2 at 5 μg/mL with PEA/FN coated scaffolds at day 7 of incubation resulted in significantly increased HBMSC ALP specific activity (p< 0.0001) compared to uncoated scaffolds in osteogenic conditions. (**Figure 2 B**). In addition, the seeding density of cells was observed to be a factor in cellular differentiation and therefore the HBMSC number seeded was increased from 2.5×10^4^ to 5×10^4^ per scaffold to ensure cellular confluence on the scaffold surface over the culture period to stimulate differentiation of the HBMSCs (**Supplementary Figure 1**). During confirmatory experiments the PEA/FN/BMP-2 coating using 100 ng/mL BMP-2 exhibited minimal ALP staining in C2C12 cells, whereas a significantly positive response was observed with a 5 µg/mL BMP-2 coating solution, indicating the BMP-2 was bound to the well/scaffold by the PEA/FN (**Supplementary Figures 2 and 3**). Furthermore, it was found that the pH of the diluent used affected BMP-2 activity (**Supplementary Figures 2, 3 and 4**).

**Figure 2:**
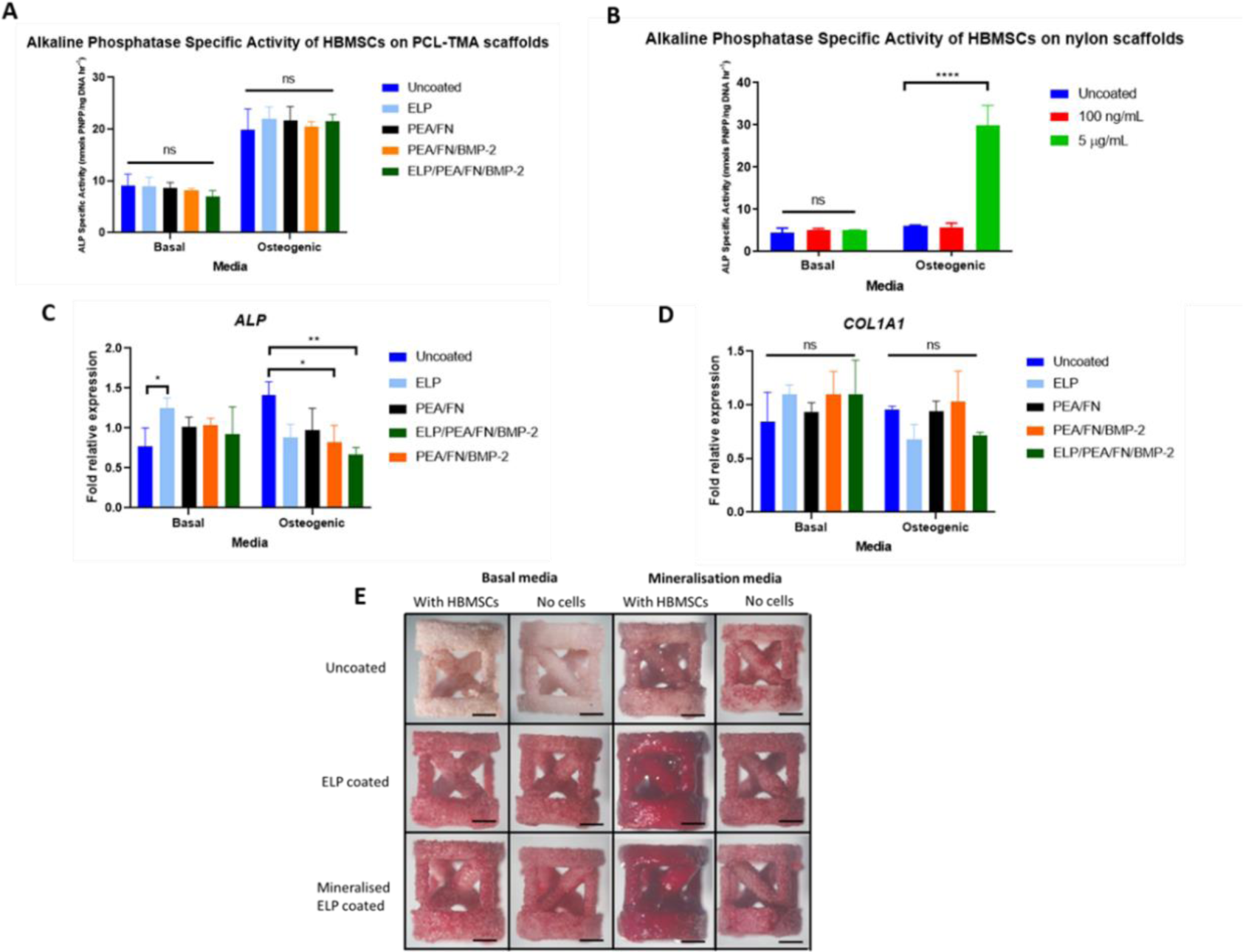
Assessment of HBMSC differentiation on coated 3D scaffold materials. (A) ALP specific activity of HBMSCs on ELP, PEA/FN/BMP-2 and ELP/PEA/FN/BMP-2 coated PCL-TMA scaffolds at day 7. There was no significant difference between the uncoated PCL-TMA and the coatings of interest in basal or osteogenic conditions. (B) ALP specific activity results comparing 100 ng/mL and 5 µg/mL BMP-2 coating solution concentration at day 7. There was a significant increase in ALP production by HBMSCs in response to the 5 μg/mL BMP-2 coating solution compared to the 100 ng/mL concentration BMP-2 solution in osteogenic culture conditions only. (C) ALP gene expression at day 7 was significantly enhanced for ELP coating than uncoated nylon in basal culture conditions, with the PEA/FN/BMP-2, ELP/PEA/FN/BMP-2 coatings displaying significantly reduced ALP gene expression in osteogenic conditions. (D) Collagen1A1 gene expression at day 7 was not significantly greater than uncoated nylon for any of the coatings in basal or osteogenic media conditions. 2-way ANOVA with Dunnett’s multiple comparisons test, n=3, mean and S.D. shown, ns; non-significant, *p<0.05, **p<0.01, ****p=<0.0001. (E) Alizarin red staining at day 27 indicated red stain uptake in the ELP coated and mineralised scaffolds when no cells were seeded, due to the constituents of the coatings themselves. ELP coating alone and with subsequent mineralisation, lead to enhanced staining due to mineral deposition on the surface of the scaffold in osteogenic culture conditions. Representative images shown, n=3, scale bar 1 mm.

The osteogenic gene expression of *ALP* and *COL1A1* were assessed for the ELP coating, PEA/FN/BMP-2 coating and combined coatings at day 14 of culture in basal and osteogenic conditions. *ALP* gene expression was significantly greater for ELP coating than uncoated PCL-TMA in basal culture conditions. In contrast, PEA/FN/BMP-2 and ELP/PEA/FN/BMP-2 coatings displayed significantly reduced *ALP* gene expression under osteogenic culture conditions (**Figure 2 C**). *COL1A1* gene expression was not significantly different under any bioactive coatings examined in comparison to uncoated PCL-TMA following culture in basal or osteogenic media conditions (**Figure 2 D**).

The ELP coated nylon scaffolds were tested in mineralisation media to determine if the HBMSCs could produce a mineralised coating on the scaffold, which was comparable to the matrix on nylon scaffolds which were mineralised in the laboratory during the coating process. ELP coated nylon scaffolds were observed to support mineralisation of matrix produced by HBMSCs following culture in an osteogenic environment, evidenced by the dark red staining, comparable to the mineralised scaffold following culture in an osteogenic environment (**Figure 2 E**). The hypothesis that a dual coating of ELP/PEA/FN/BMP-2 would be synergistic *in vivo* compared to each coating alone was examined. The assumption was that HBMSCs would adhere to the scaffolds initially, due to the presence of FN and BMP-2 followed by enhanced mineralised matrix in response to the ELP/BMP-2. Therefore, the ELP, PEA/FN/BMP-2 and ELP/PEA/FN/BMP-2 coating options on PCL-TMA scaffolds were assessed in the CAM assay.

### 3.2 The PCL-TMA material and bioactive scaffolds were biocompatible following evaluation on the CAM assay

PCL-TMA scaffolds with ELP, PEA/FN/BMP-2 and ELP/PEA/FN/BMP-2 coatings were analysed for biocompatibility and angiogenic support using the CAM assay. All three coatings on PCL-TMA showed excellent biocompatibility and the ability to support angiogenesis, although viability scores were reduced due to undeveloped chicks (**Figure 3 A**). Analysis demonstrated no significant difference in Chalkley score between uncoated PCL-TMA scaffolds and PCL-TMA scaffolds with coatings, when analysed by a blinded observer (**Figure 3 B**). The scaffolds were all noted to integrate with the CAM with blood vessels clearly visible, surrounding each scaffold, with no gross thickening or disruption of the CAM tissue due to the scaffolds (**Figure 3 C**). The PCL-TMA scaffolds were noted to be surrounded by CAM tissue, blood vessels and host cells from the CAM evidenced following Alcian blue and Sirius red or Goldner’s trichrome staining (**Figure 3 D**). The brittle nature of the PCL-TMA led to the polymer material shearing on sectioning, despite being embedded in hard GMA resin, potentially indicating limitations to this PCL-TMA material. Overall, the various bioactive coatings were biocompatible when combined with the PCL-TMA material, with no indication of an inflammatory response observed.

**Figure 3:**
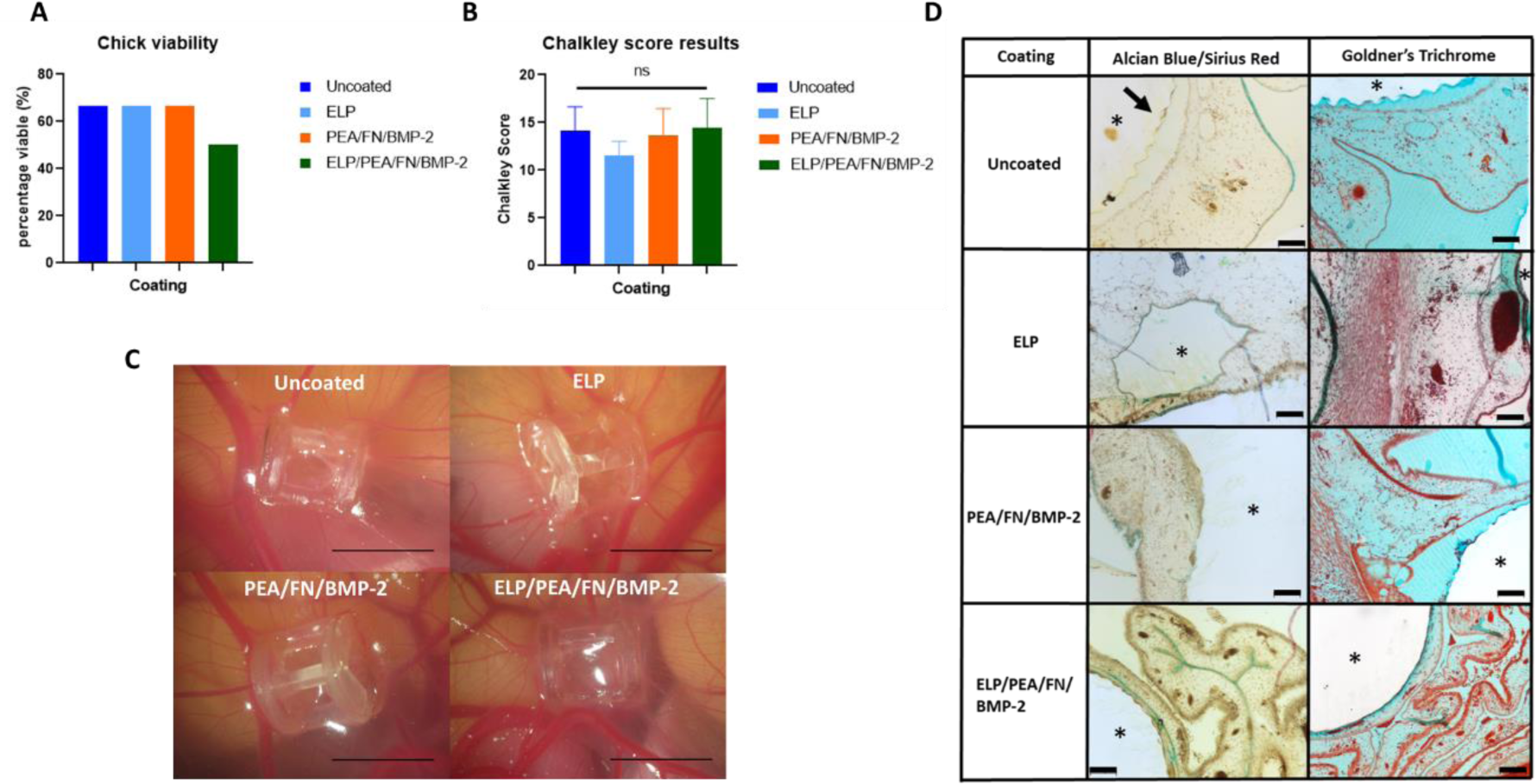
CAM assay viability and Chalkley score results for PCL-TMA scaffolds. (A) Chick viability was suboptimal due to poor chick development, n=6. (B) There was no significant difference in Chalkley score between uncoated PCL-TMA and the coated scaffolds, (uncoated n=4, ELP n=4, PEA/FN/BMP-2 n=4, ELP/PEA/FN/BMP-2 n=3), ns=non-significant. One-way ANOVA with Dunnett’s multiple comparisons test was used for statistical analysis, mean and S.D. shown. (C) Photographs of representative uncoated PCL-TMA and ELP coating, PEA/FN/BMP-2 and ELP/PEA/FN/BMP-2 coated PCL-TMA scaffolds on the CAM. The PCL-TMA material, the ELP and PEA/FN/BMP-2 coatings were biocompatible and supported angiogenesis. Scale bar 5 mm. (D) Histological staining (Alcian blue and Sirius red or Goldner’s trichrome) of PCL-TMA scaffolds surrounded by CAM tissue. The PCL-TMA scaffold material did not support sectioning, with fragments remaining (arrow), but tissue around the prior scaffold (*) could be determined. Scale bar 100 μm.

### 3.3 Bone formation analysis on the PCL-TMA scaffolds compared to collagen sponge with BMP-2 using the mouse subcutaneous implantation assay

#### 3.3.1 µCT analysis results of uncoated scaffolds compared to coated PCL-TMA scaffolds and collagen sponge/BMP-2

The mouse subcutaneous implantation study demonstrated that the PCL-TMA scaffolds displayed no detectable mineralisation at week 2, 4 or 6 scans *in vivo*. In contrast, extensive mineralisation was observed in the collagen sponge with 5 µg BMP-2 from week 2 onwards, clearly seen in representative week 6 images, as a mineralised disc (**Figure 4 A**). The excised samples from the mice were scanned *ex vivo* at week 8 and no significant mineral was detected in response to either of the ELP or PEA/FN/BMP-2 coatings, or when applied concurrently. The collagen sponge displayed significant mineralisation due to the response of the adipose and subcutaneous stromal cells to the BMP-2, when quantified at week 8, compared to the uncoated PCL scaffolds (**Figure 4 B (i)**). There was no significant difference between the negligible quantities of bone formed on the coated scaffolds compared to the uncoated PCL-TMA scaffolds (**Figure 4 B (ii)**).

**Figure 4:**
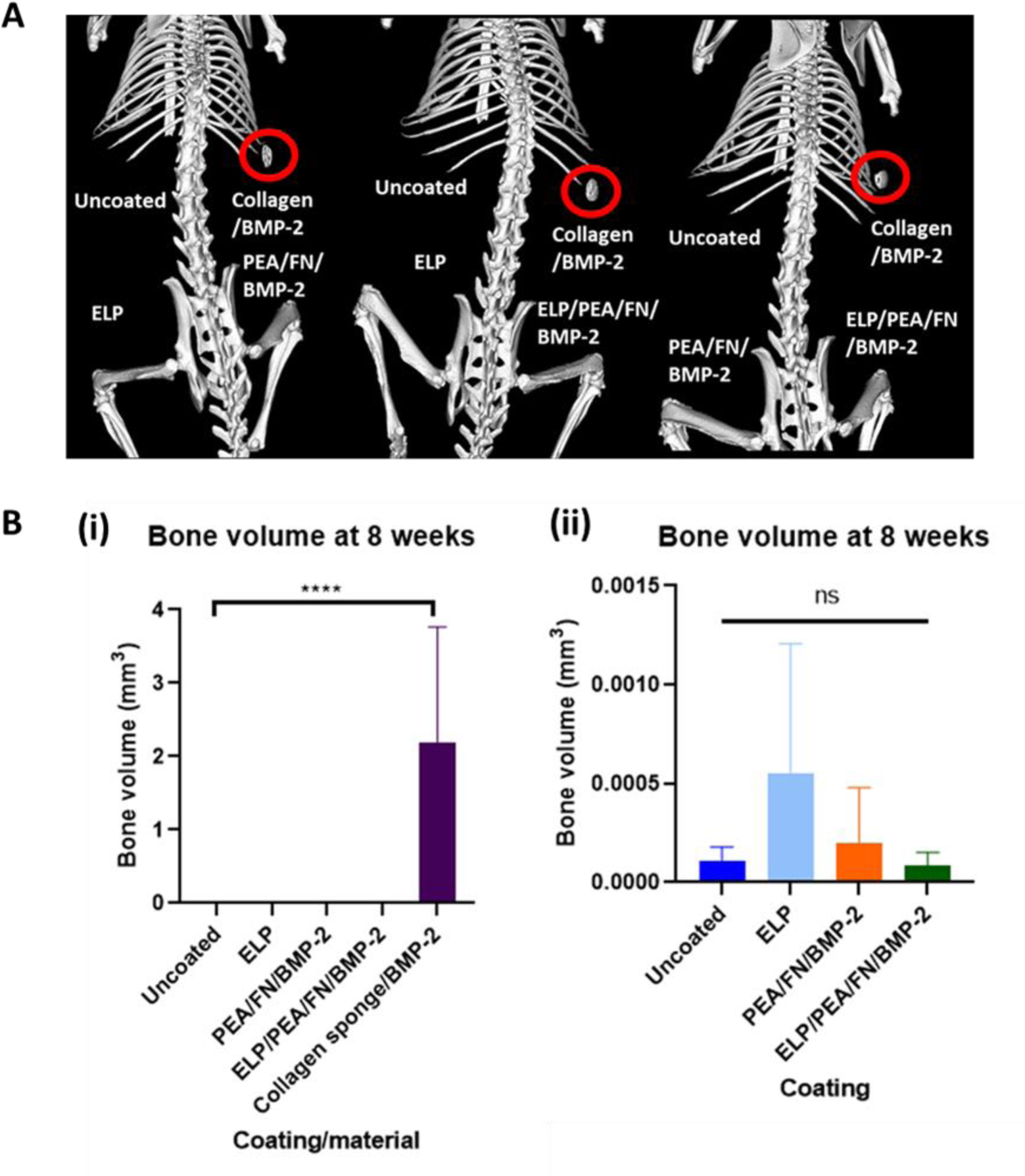
µCT results of the murine subcutaneous implantation study. (A) Representative µCT images with no bone formation observed in uncoated, PEA/FN/BMP-2, ELP, or ELP/PEA/FN/BMP-2 coated PCL-TMA scaffolds, however the collagen sponge with 5 µg of BMP-2 showed mineralisation in all mice (within red circles) at week 6. (B) Quantification of bone volume formed in the mouse subcutaneous implant model using PCL-TMA scaffolds and collagen sponge/BMP-2. (i) The collagen sponge displayed significant bone formation at 8 weeks compared to the uncoated PCL-TMA scaffold. (ii) There was no significant difference between the negligible bone formation on the coated scaffolds compared to the uncoated PCL-TMA scaffolds. One-way ANOVA with Dunnett’s multiple comparisons test, ns= not significant, ****p<0.0001. N=9 uncoated scaffolds, n=9 collagen sponge, n=6 PEA/FN/BMP-2 and n=6 ELP/PEA/FN/BMP-2 coated scaffolds, mean and S.D. shown.

#### 3.3.2 Histological analysis of the subcutaneous PCL-TMA scaffolds and positive control collagen sponge/BMP-2

Histological analysis of the PCL-TMA bioactive scaffolds following subcutaneous implant studies showed discrete shards of fragmented PCL-TMA material from the periphery of each PCL-TMA scaffold (coated and uncoated) (example **Figure 5 A**). The surrounding tissue was integrated with the scaffold as evidenced by the scaffold ridges within the tissue (example **Figure 5 J**). Alcian blue and Sirius red staining indicated the presence of proteoglycans within the surrounding tissue and a discrete red rim of bone tissue staining at the perimeter of the collagen sponge with BMP-2 (**Figure 5 F**). Goldner’s trichrome stain demonstrated collagenous bone matrix (green stain) in the collagen sponge/BMP-2 group (**Figure 5 F**), while the muscle tissue was stained vivid red in the uncoated scaffold example (**Figure 5 B**). Intensely red staining collagenous tissue was observed to surround the collagen sponge and the PEA/FN/BMP-2 coated scaffold (**Figures 5 F and N**). Alizarin red staining produced a marked dark red stain around the collagen sponge with BMP-2, indicative of mineralisation as seen on µCT analysis (**Figure 5 G**). No detectable bone formation was seen on histological examination of the coated or uncoated PCL-TMA scaffolds (**Figures 5 A - D, I - T**). Von Kossa staining was comparable in location to the alizarin red, confirming the presence of mineralisation around the periphery of the collagen sponge/BMP-2 construct only (**Figure 5 H**). The perimeter of all PCL-TMA scaffolds examined showed no positive black staining, confirming an absence of bone formation (**Figures 5 D, P, L, T**).

**Figure 5:**
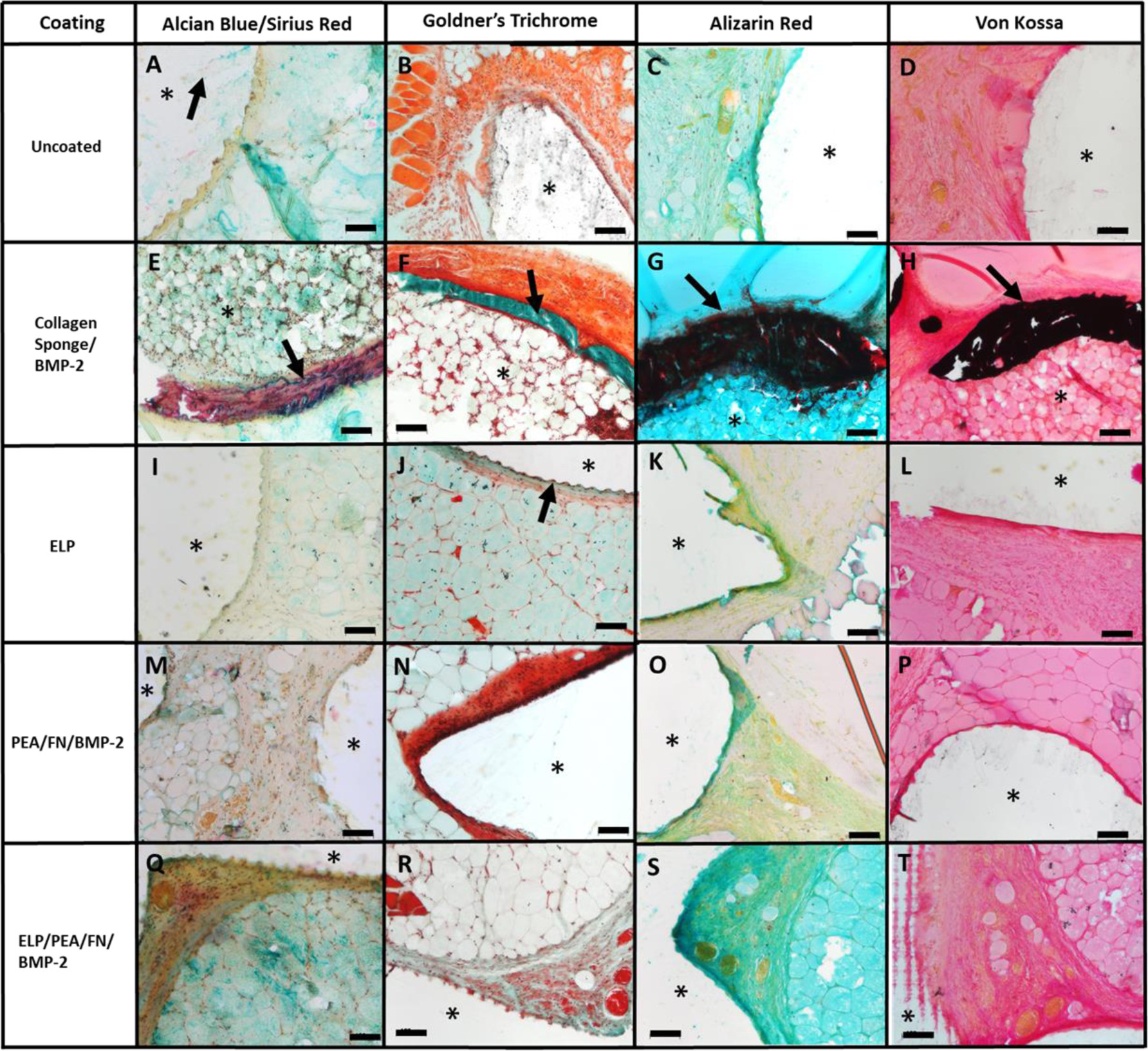
Alcian blue and Sirius red, Goldner’s trichrome, Alizarin red and Von Kossa staining of PCL-TMA scaffolds and collagen sponge/BMP-2. The scaffolds were not amenable to sectioning; however, the surrounding tissue remained. (*) PCL-TMA scaffold area or collagen sponge/BMP-2. (A) Shards of PCL-TMA material (black arrow) remain in the section. (B) Vivid red staining muscle was seen but no bone formation. (C and D) No bone formation was found on the uncoated scaffold. (E - H) Only the collagen sponge displayed marked mineralisation and bone formation around the periphery (arrows). (I - T) No bone formation was seen on the ELP, PEA/FN/BMP-2 or ELP/PEA/FN/BMP-2 coated scaffolds, with the ridges of the scaffold material seen surrounded by tissue (J arrow). Scale bar 100 μm.

## 4. Discussion

The current study set out to examine the potential of bioactive 3D-printed octetruss PCL-TMA scaffolds for bone formation augmentation. The *in vitro* studies examined cytocompatibility, the ability to induce ALP and mineral production and to enhance osteogenic gene expression within HBMSCs. All of the bioactive coatings examined showed no signs of cytotoxicity after 14 days and viable cells were visualised with live and dead cell labelling. CAM assay analysis demonstrated biocompatibility of the PCL-TMA scaffold material and bioactive coatings with no effect on angiogenesis or chick development. The subcutaneous implantation study revealed marked bone formation around the collagen sponge with BMP-2, while there was no bone formation on any of the bioactive coated PCL-TMA scaffolds.

Several factors are known to affect cell differentiation including: i) material stiffness, ii) hydrophobicity, iii) the concentration, configuration and subsequent activity of growth factor due to the method of application and, iv) the culture time may impact on cell differentiation. The ALP specific activity results were not significantly different from uncoated scaffolds in basal media conditions for the PEA/FN/BMP-2 coating, with the bound BMP-2 unable to induce significant osteogenic differentiation of cells. It has been shown that less than 10% of the bound BMP-2 is released over a 14-day period, thus to explain the *in vitro* and *in vivo* findings, if only a small quantity is bound due to the relatively low surface area of the PCL-TMA material or possible hydrophobicity impacting on PEA coating uniformity, this may have led to the PEA/FN/BMP-2 coating being less efficacious than in reported studies [13]. Therefore, based on this finding it was difficult to make a firm conclusion given that the *in vitro* environment with encouraging results differs from the *in vivo* environment, limiting assessment of the coating capabilities. Interestingly, the PEA/FN/BMP-2 coating using 100 ng/mL BMP-2 exhibited minimal ALP staining in C2C12 cells, whereas a significantly positive response was observed with a 5 µg/mL BMP-2 coating solution on tissue culture plastic and PCL circular scaffolds, indicating the BMP-2 was bound to the well/scaffold by the PEA/FN. However, this PCL scaffold was of a different material type, hydrophobicity, and surface area to the PCL-TMA scaffolds and thus may permit greater PEA/FN/BMP-2 adhesion capability, highlighting the importance of optimisation of the underlying scaffold material.

Different species have differing sensitivity to BMPs, with human osteoblasts requiring dexamethasone with BMP-2, 4 or 7 to increase ALP activity compared to murine osteoblasts [22-24]. This effect was potentially observed in the current study, when HBMSCs were cultured on the PEA/FN/BMP-2 (5 μg/mL) coated nylon scaffolds in osteogenic media. This higher concentration of 5 μg/mL BMP-2 was used in the CAM assay and murine *in vivo* studies, however, *in vitro* cell culture experiments do not always replicate *in vivo* cell differentiation [25]. Interestingly, the findings at 5 μg/mL in basal conditions *in vitro* may mirror our *in vivo* findings in a heterotopic site. However, the ability of the low mass of BMP-2 bound to scaffolds for prolonged periods of time to induce osteogenic differentiation of HBMSCs in osteogenic conditions *in vitro* was encouraging. This indicates potential for osteogenesis in an osteogenic *in vivo* environment, therefore using a bone defect model may be more appropriate.

As detailed in the supplementary information, an initial coating was undertaken using acidic buffer solution (pH 4.5) to dilute the InductOs® rhBMP-2, as pH was thought to be inconsequential on the success of the coating method or cell differentiation. However, the current studies indicate the importance of the pH employed. It is reported that InductOs® BMP-2 activity is pH dependent and aggregation occurs at pH 6.5, hence in order to prevent this, the buffer was established at pH 4.5 [26]. However, when testing bioavailability and activity of BMP-2 on C2C12 cells diluted in the acidic buffer, BMP-2 was found to be either inactive or unbound to the FN. A neutral or basic environment can destabilize the disulfide bonds, damaging the dimer configuration and inactivating the BMP, however InductOs® BMP-2 generated a positive ALP staining in the neutral buffer at a higher concentration of 5 μg/mL [27].

The difference in ALP gene expression when the ELP coating was applied alone compared to application of PEA/FN/BMP-2 on top of the ELP coating, may result in the PEA/FN/BMP-2 coating prohibiting the action of the ELP coating. It is known that gene expression does not always relate to the production of matrix or protein at the scaffold surface or implant/bone interface, therefore the higher ALP gene expression in the ELP coating only group did not appear to translate to higher ALP protein formed in ALP specific activity assessment [28]. The ELP coating on nylon scaffolds was found to have the potential to be mineralised by HBMSCs when exposed to a mineralising media environment visualized by alizarin red staining. Thus, the PCL-TMA scaffolds with ELP coating and/or PEA/FN/BMP-2 coatings were found to be cytocompatible and support osteogenic differentiation and mineralisation of the microenvironment, therefore these bioactive coating options were further assessed *in vivo*. The CAM assay provided an intermediary step with results for the uncoated PCL-TMA, ELP coating, PEA/FN/BMP-2 coating and the combined ELP/PEA/FN/BMP-2 coatings demonstrating excellent biocompatibility and angiogenic response using accepted angiogenic quantitation methodology [29-32].

The mouse subcutaneous implantation model allowed the first stage of scaffold assessment in a rodent preclinical model and confirmed the biocompatibility of the PCL-TMA scaffold and coatings. The study also confirmed the inability to mineralise in a heterotopic site, compared to the commercially available and clinically applied collagen sponge with BMP-2. Bone did not form on the scaffold constructs, whereas collagen sponge with BMP-2 displayed significant bone formation, highlighting the need for consideration of effective growth factor mass, as the mass of BMP-2 (5 µg) within the collagen sponge was likely much greater than that attached to the PCL-TMA scaffolds and need for appropriate animal model use. The objective was to determine the optimal osteogenic coating on the PCL-TMA scaffold; however, the results did not determine one optimal coating from the ELP and/or PEA/FN/BMP-2 options using the subcutaneous implantation model, despite a sufficient period for mineralisation to occur. The PEA/FN/BMP-2 coating has been used successfully in osseous defect sites, with as little as 15 ng of BMP-2 in the murine radial defect model [13]. An ELISA using BMP-2 solutions to determine the mass of BMP-2 adhered to the PEA/FN coated PCL-TMA scaffolds was not performed due to difficulties attributed to aggregation of the protein following freezing at -20 °C [33]. It may be that the quantity of BMP-2 bound from a 5 µg/mL solution to the PCL-TMA material may be less than required for a response by the subcutaneous tissues given the ‘critical mass’ required to induce mineralisation in rodents has been indicated to be around 1 µg of BMP-2, from review of the published effective and ineffective doses used [34]. A challenge is the lack of response that human skeletal cell populations display, assessed via ALP activity, to BMP-2 compared to rodent derived cells which can lead to discrepancy between rodent models and human clinical data [24]. Reporting the concentration and volume of growth factor use in animal models and details of material size and surface area allows more accurate comparison between animal studies, especially subcutaneous implant studies, as these vary as reported by Gothard *et al* [35]. Consideration for the coating method and mass of growth factor delivered, often limited by the surface area of the scaffold, possibly limit the efficacy of this PEA/FN/BMP-2 coating when used in this model. Therefore, an osseous defect site may enhance the response to BMP-2 presentation by the scaffold or if the scaffold is optimised to increase the BMP-2 binding by altering the surface area and hydrophobicity for PEA/FN coating. The theoretical action of the ELP coating entails its ability to mineralise by sequestering surrounding calcium and phosphate ions, however, it is possible that the ELP coating could not access mineral to ‘grow’ on the surface of the ELP coating in the subcutaneous space. Therefore, application in an osseous defect model may offer a more suitable model and environment to test this bioactive coating.

## 5. Conclusion

The PCL-TMA material and bioactive coatings were cytocompatible and the ELP coating and PEA/FN/BMP-2 coated scaffolds supported osteogenic differentiation of HBMSCs. The ELP coating enhanced the osteogenic ALP gene response of HBMSCs in basal conditions on PCL-TMA scaffolds and permitted mineralisation of the ELP coating by HBMSCs in mineralising osteogenic media conditions. The PEA/FN/BMP-2 coated scaffolds showed marked ALP production by C2C12 cells when used at the *in vivo* concentration (5 µg/mL) and in osteogenic conditions with HBMSCs. The CAM assay demonstrated the PCL-TMA scaffold material and bioactive coatings of ELP coating, PEA/FN/BMP-2 and ELP/PEA/FN/BMP-2 were supportive of angiogenesis and confirmed their biocompatibility. However, the ELP, PEA/FN/BMP-2 and ELP/PEA/FN/BMP-2 coatings were ineffective in inducing bone formation in the subcutaneous implantation model of heterotopic bone formation. Despite *in vitro* data suggesting the scaffolds could induce bone formation, the coatings did not mineralise in an *in vivo* subcutaneous environment. These results indicate the importance of the selection of an appropriate model for material/growth factor assessment and bone formation mechanism determination. An osseous site defect may be more appropriate to test the ELP, PEA/FN/BMP-2 and ELP/PEA/FN/BMP-2 coatings if a more vascular, osteogenic site is required to initiate or enhance the response of skeletal cells to these coatings. Therefore, prior to discounting these coating materials for clinical translation, the use of a rodent bone defect model would be warranted.

### Credit author statement

Øvrebo, Ø. designed the scaffold shape using C.A.D software, Echalier, C. and Wojciechowski, J. P. synthesised the PCL-TMA resin, Wojciechowski, J. P., Echalier, C. and Yang, T. prepared and printed the PCL-TMA scaffolds, Jayawarna, V. printed the circular PCL scaffolds, performed the PEA coating and organised the EO sterilisation of materials, Hasan, A. performed the ELP coating of scaffolds, Marshall K. M. conducted the *in vitro* experiments, the CAM assay, murine subcutaneous implantation study, data collection and analysis, histology and wrote the paper. Kanczler J. M., Mata, A., Salmeron-Sanchez, M., Stevens, M. M. and Oreffo R. O. C. are credited for conceptualisation, funding acquisition, study supervision and editing of the manuscript.

### Data and materials availability

All data associated with this study are presented in the paper or the Supplementary Materials. All raw data is available on reasonable request from the corresponding authors.

## Supporting information

Supplemental information Bioactive coatings on PCL scaffolds

## Acknowledgements

Research support for this study from the UK Regenerative Medicine Platform Acellular / Smart Materials – 3D Architecture (MR/R015651/1), the Biotechnology and Biological Sciences Research Council (BBSRC BB/P017711/1), ERC Proof-of-concept grant MINGRAFT and the University of Southampton is gratefully acknowledged as well as many useful discussions with past and current members of the Bone and Joint Research Group in Southampton, UK. Dr Katie Dexter (University of Southampton, Biomedical Imaging Unit) is acknowledged for advice and expertise in µCT scanning of mice. Jon Ward (University of Southampton, Histochemistry Research Unit) is acknowledged for his teaching of histological methods. We would like to thank Robyn Andrews, Dr Susannah Clarke and Embody Orthopaedics for 3D printing the Nylon scaffolds. Dr Akemi Nogiwa Valdez is acknowledged for proof reading of the manuscript. For the purpose of open access, the author has applied a ‘Creative Commons Attribution (CC BY) license to any Author Accepted Manuscript version arising.

## Appendix A. Supplementary data

Supplementary data to this article can be found online.

