## Supplemental information Bioactive coatings on PCL scaffolds for "Bioactive coatings on 3D printed scaffolds for bone regeneration: Translation from *in vitro* to *in vivo* models and the impact of material properties and growth factor concentration"

### Supplementary information

#### Low cell density limits ALP production by HBMSCs in osteogenic culture conditions

Due to unexpected ALP specific activity results, the cell seeding number was investigated to optimise HBMSC differentiation in osteogenic culture conditions for biochemistry and molecular gene expression experiments. A seeding density of $5x10^4$, $1x10^5$ and $2x10^5$ HBMSCs led to a significant difference between ALP specific activity in basal and osteogenic culture conditions, therefore $5x10^4$ HBMSCs was chosen for ALP specific activity and molecular experiments (**Supplementary Figure 1**).

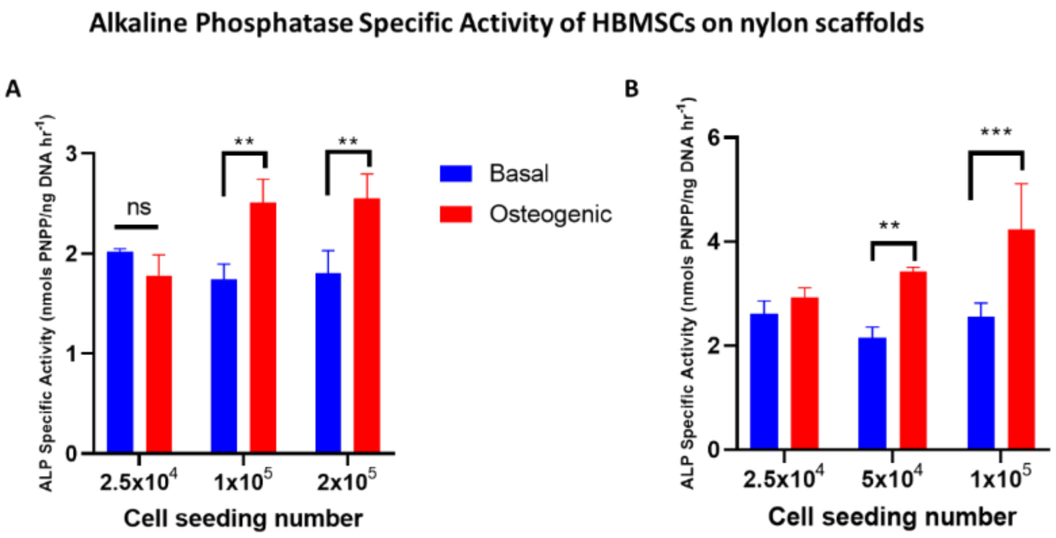


**S. Figure 1:** The effect of cell seeding number on ALP specific activity. The greater cell seeding number led to a significant difference in ALP specific activity with (A) $1x10^5$ and $2x10^5$ and (B) $5x10^4$ and $1x10^5$ HBMSCs displaying a significant difference between basal and osteogenic culture conditions compared to $2.5x10^4$ HBMSCs which was initially used. N=3, 2-way ANOVA with Šidáks multiple comparisons test, mean and S.D, ns; non-significant, $^{**}p<0.01$, $^{***}p<0.001$.

#### *BMP-2 acidic buffer solution constituents*

The buffer solution for dilution of Medtronic InductOs® BMP-2 was made by dissolving the following listed in Supplementary Table 1 in deionised water:

**S. Table 1:** InductOs® BMP-2 buffer solution constituents.

| **Reagent** | **Concentration** | **Quantity to make 500 mL** |
|---|---|---|
| Sucrose | 0.5% | 2500 mg |
| Glycine | 2.5% | 12500 mg |
| L-glutamic acid | 0.37% | 1850 mg |
| Sodium chloride | 0.01% | 50 mg |
| Polysorbate 80 | 0.01% | 50 mg |

*Alkaline phosphatase analysis for assessment of osteogenic differentiation*

At day 7, cells on TCP or scaffolds in 24 well plates were washed in PBS twice and fixed in 90% ethanol for 10 minutes, prior to washing again in PBS. Alkaline phosphatase (ALP) activity was measured by creating a solution of 4% (v/v) Naphthol AS-MX phosphate and 0.0024% (w/v) Fast Violet-B salt mixed in distilled water and adding 300 µL per well. Cells were incubated in the dark in the incubator at 37 °C for 30-60 minutes. Deionised water was added to stop the reaction and the solution removed prior to imaging. A red/purple colour indicated a positive result of ALP activity. A Canon G10 camera attached to a stemi 2000-c stereomicroscope (Zeiss, UK) was used to record images, taken using the same magnification and camera brightness settings to allow subjective comparisons of staining intensity to be made.

*Experimental method for the analysis of the effect of diluent pH on the bioactivity of BMP-2*

To verify the ability of the PEA/FN/BMP-2 coating to differentiate C2C12 cells to an osteogenic phenotype, the BMP-2 was prepared in acidic buffer of pH 4, or neutral pH 7 PBS. UV sterilised, PEA coated 24 well plates had FN and BMP-2 coating to analyse osteogenic differentiation of C2C12 cells (ATCC (CRL-1772)), verified by ALP staining. PEA/FN were used to coat the negative control wells and conditions were in triplicate. The wells to test the acidic buffer diluent were rinsed with PBS prior to addition of 300 µL FN/PBS with $CaCl_2$/$MgCl_2$ (20 µg/mL) for 1 hour at RT on the horizontal plate shaker at 180 rpm. The FN solution was removed and the wells rinsed with PBS. A 1 mL aliquot of InductOs® BMP-2/acidic buffer (5 µg/mL or 100 ng/mL) was subsequently added and incubated at RT for 1 hour on the horizontal plate shaker at 180 rpm. The BMP-2 solution was removed and the wells rinsed in PBS and allowed to dry. Alternatively, wells to test the neutral diluent of PBS with $CaCl_2$/$MgCl_2$ were rinsed with and had the FN and BMP-2 diluted in PBS with $CaCl_2$/$MgCl_2$. $1\times10^5$ C2C12 P29 cells in 1 mL of media were added to each well and cultured at 37 °C in 5% $CO_2$/balanced air. Media consisted of DMEM/1% P/S with or without BMP-2 (100 ng/mL positive control) and 10% FCS was added to each well after 3 hours of culture. Media was changed to DMEM/1% P/S/2% FCS after 24 hours (day 1) and on day 4. ALP staining was performed on day 7 and photographs captured using a stemi 2000-c stereomicroscope and Canon G10 digital camera.

*Effect of diluent pH on C2C12 cell differentiation in a coated well plate*

The acidic buffer showed no ALP staining at 5 µg/mL or 100 ng/mL BMP-2 concentration, indicating the C2C12 cells did not differentiate. In contrast, 5 µg/mL BMP-2 diluted in PBS (with $CaCl_2$ and $MgCl_2$ added) did induce osteogenic differentiation of C2C12 cells. The positive control of BMP-2 100 ng/mL within the media gave intense ALP staining while no staining was seen in the negative control wells (**Supplementary Figure 2**).

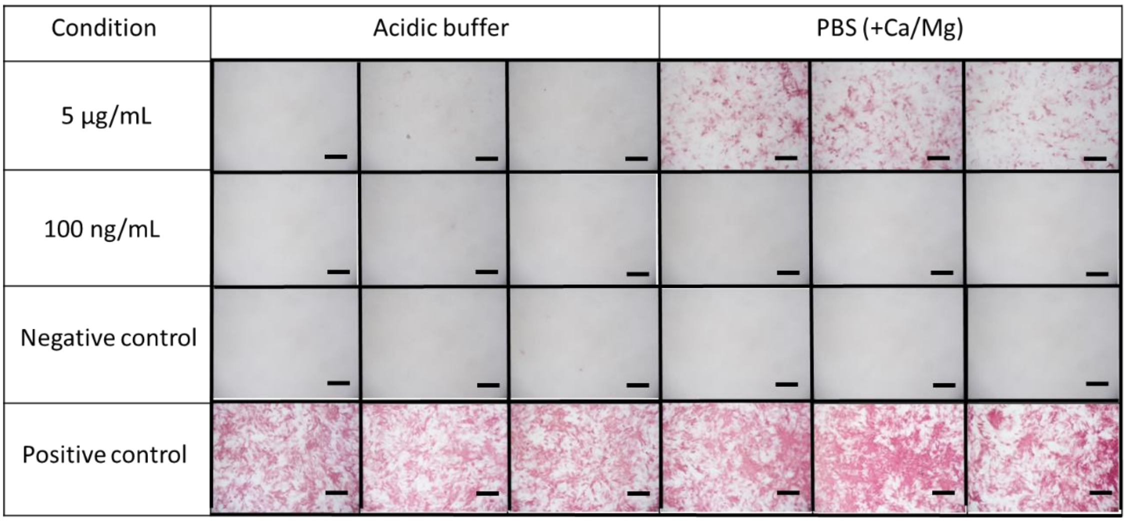


**S. Figure 2:** ALP staining of C2C12 myoblast cells cultured using coated well plates with different diluents. A 24 well plate coated using BMP-2 diluted in acidic buffer or neutral PBS showed positive red ALP staining at 5 µg/mL. No stain was visible at 100 ng/mL BMP-2 or in the negative control where BMP-2 was not used to coat the plate nor added to the media. The positive control represents BMP-2 100 ng/mL added to the culture media with a marked positive red staining response. N=3, Scale bar 500 µm.

#### *Seeding C2C12 cells onto circular PCL scaffolds to confirm the effect of diluent pH on BMP-2 activity*

This experiment confirmed the effect of BMP-2 diluent pH and verified the activity of the BMP-2 on PEA/FN coated 3D scaffolds. 3D extrusion printed, circular (5mm diameter by 0.8mm high), porous, lattice shaped PCL scaffolds were EO sterile uncoated, or PEA coated. For coating, PBS was the diluent for FN and for rinsing, with acidic buffer (Supplementary Table 1) for BMP-2 in the acidic group; or PBS containing $CaCl_2$ and $MgCl_2$ was used for FN dilution, rinsing and BMP-2 dilutions in the neutral pH group. C2C12 cells (ATCC (CRL-1772)) were seeded at $2x10^5$ cells per scaffold in 25 µL media (DMEM/1%PS/10%FCS with 100 ng/mL BMP-2 for positive controls). After 15-30 minutes 475 µL of the relevant media type was added to give 500 µL total volume. After 24 hours, the scaffolds were moved to a new 24 well plate and the media changed to include 2% FCS, which was changed every 3 days and ALP staining performed on day 7. The test scaffolds were n=3, however n=1 for controls of PEA or PEA/FN coated in the neutral pH group.

##### *Effect of diluent pH on C2C12 cell differentiation on 3D PCL scaffolds*

The 2D results were subsequently confirmed by culture of C2C12 cells on 3D PCL scaffolds with the lack of staining at 100 ng/mL in both the acidic and neutral pH diluents, but a marked positive red stain response was seen in the 5 µg/mL neutral PBS diluent group. The positive control groups showed staining, illustrating that the BMP-2 from the media was effective, but the low pH may have inhibited the BMP-2 binding to the FN in the coated group (**Supplementary Figure 3**).

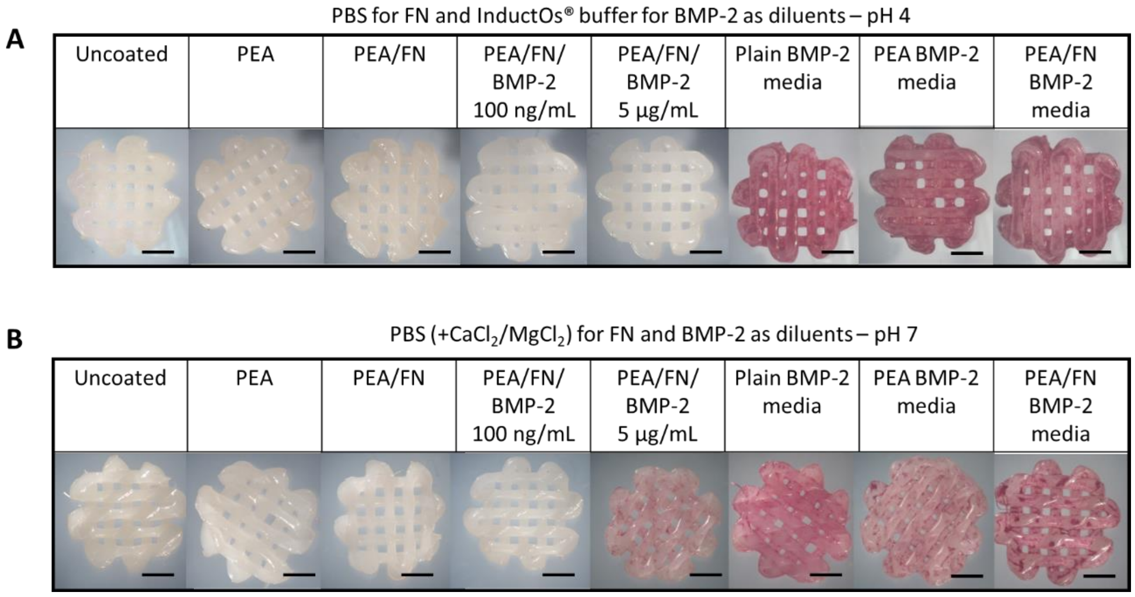


**S. Figure 3:** The effect of diluents on BMP-2 activity observed by ALP staining of C2C12 cells. Low pH does not appear to allow BMP-2 to bind to the scaffold even at high concentrations, while the neutral pH facilitates binding. 100 ng/mL was insufficient to induce differentiation of the C2C12 cells compared to 5 µg/mL BMP-2 in neutral pH conditions. Representative images shown, n=3 for all conditions except n=1 for PEA and PEA/FN in BMP-2 media in the pH 7 group. Scale bar 1 mm.

##### *Analysis of BMP-2 source to compare the efficacy of BMP-2 at low concentrations*

To assess if R&D BMP-2 was more efficacious at a low concentration of 100 ng/mL than InductOs® BMP-2, PEA coated 24 well plates were used to adhere FN and BMP-2 (from Medtronic, USA, or R&D, Biotechne, UK) for analysis of the osteogenic differentiation of C2C12 cells. The wells of a UV sterile 24 well plate were rinsed with PBS with $CaCl_2/MgCl_2$ and allowed to dry. A 300 µL aliquot of FN/PBS with $CaCl_2/MgCl_2$ solution (20 µg/mL) was added for 1 hour at RT on the horizontal plate shaker at 180 rpm, removed and rinsed with PBS with $CaCl_2/MgCl_2$. PBS with $CaCl_2/MgCl_2$ was the diluent for both types of BMP-2 as it was shown to maintain bioactivity of BMP-2. InductOs® BMP-2 (5 µg/mL, 100 ng/mL and 50 ng/mL) and R&D BMP-2 (100 ng/mL and 50 ng/mL), 1 mL per well in triplicate was incubated 1 hour at RT on the horizontal plate shaker at 180 rpm. The BMP-2 solution was removed and the wells rinsed in PBS with $CaCl_2/MgCl_2$ and allowed to dry. Media consisted of DMEM/1% P/S ± BMP-2 (100 ng/mL R&D or InductOs® BMP-2) and 10% FCS was added to each well after 3 hours of culture. $1\times10^5$ C2C12 P29 cells in 1 mL of appropriate media were added to each well and cultured at 37 °C in 5% $CO_2$/balanced air. Media was changed to DMEM/1% P/S/2% FCS after 24 hours (day 1) and on day 4. ALP staining was performed on day 7.

*Variation in BMP manufacturer on bioactivity of BMP-2*

The assay was performed using the neutral buffer with R&D BMP-2 (R&D systems, Biotechne) and InductOs® BMP-2 (Medtronic, USA) and neither BMP-2 type induced osteogenic differentiation of C2C12 cells at 100 ng/mL or 50 ng/mL BMP-2 concentration. A 5 µg/mL solution of InductOs® BMP-2 showed ALP staining again, confirming the reproducibility of the process when InductOs® BMP-2 was used (**Supplementary Figure 4**).

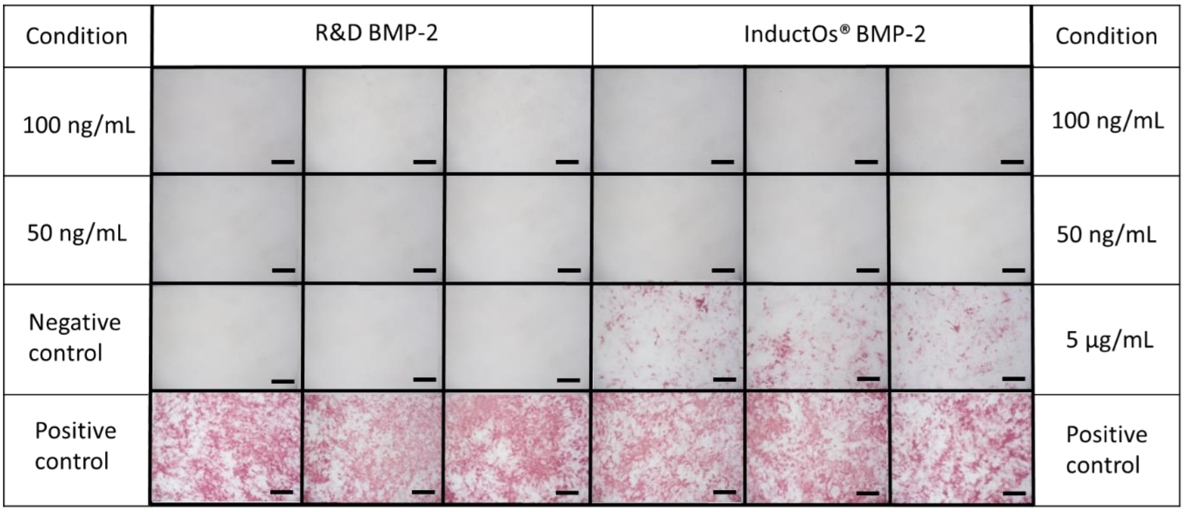


**S. Figure 4:** ALP staining of C2C12 myoblast cells after exposure to BMP-2 from different suppliers. A 24 well plate coated using R&D or InductOs® BMP-2 in PBS (with $CaCl_2$/$MgCl_2$) shows positive red ALP staining at 5 µg/mL of InductOs® BMP-2 which confirmed previous findings. The negative control had no visible staining indicating no spontaneous ALP production. 100 ng/mL and 50 ng/mL R&D and InductOs® BMP-2 were both negative with no ALP stain visible. The positive controls of both R&D and InductOs® BMP-2 added to the media directly shows intense red ALP staining indicative of ALP production by C2C12 cells. Scale bar 500 µm, n=3 for each condition.

#### Method of albumin removal from the eggs for the CAM assay

Albumin removal facilitated 'dropping' the CAM away from the inner shell membrane to prevent tearing of the CAM during windowing and reduced scaffold entrapment between the eggshell and CAM, hampering blood vessel quantification by the Chalkley scoring method. At ED 3, the egg was candled and a cross drawn on the upper surface at the embryo to ensure orientation (**Supplementary Figure 5 A**). A sterile Schedule II Laminar flow cabinet with a sterile egg box, sterile disposable no 10-scalpel blade/handle, 5 mL syringes, 19-gauge 2-inch needles and 23-gauge 1-inch needles, Parafilm squares (2 cm$^2$) in 70% ethanol and autoclave tape to secure the parafilm was set up. The pointed end of the egg was wiped with Chemgene disinfectant (Medimark Scientific). A hole was made by twirling the blade round repeatedly (**Supplementary Figure 5 B**). Eggshell dust confirmed progress and this process became faster, as abrasive dust accumulated on the blade. A 19-gauge needle was inserted horizontally into the hole, to the bevel (about 5mm) (**Supplementary Figure 5 C**).

A 23-gauge needle attached to a 5 mL syringe, with the increments visible to the user, was used to withdraw 3 mL of albumin (**Supplementary Figure 5 D**). Insertion of the needle in a downwards direction with the bevel of the needle uppermost aided albumin retrieval, with constant steady pressure applied on the syringe to avoid disturbing the yolk, creating froth or blockage of the needle from thick albumin. The hole was covered with Parafilm (soaked in 70% ethanol, rinsed in PBS and drip dry) and autoclave tape (**Supplementary Figure 5 E and F**). A second piece of tape about 50% below the first piece created an overhanging tape edge, so the egg would reside horizontally rather than tipping in the incubator, if necessary. The eggs were placed horizontally within an egg incubator at 37 °C and 60% humidity without rotation.

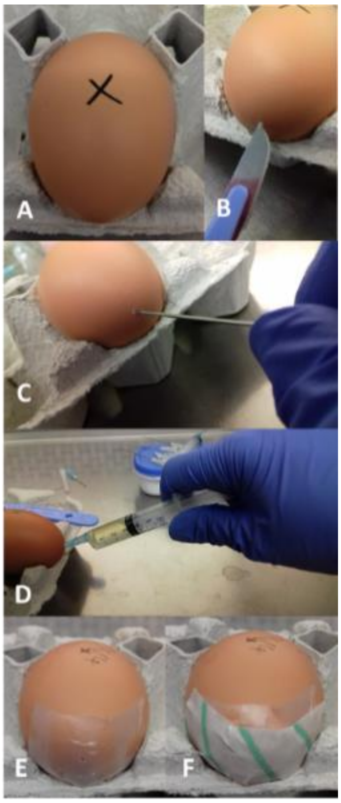


**S. Figure 5:** Albumin removal from eggs at ED 3 for the CAM assay. (A) A cross was drawn on top of the egg where the embryo was to ensure the egg was kept upright. (B) A hole in the pointed end of the eggshell was made using a scalpel blade. (C) A needle was used to make a hole in the inner shell membrane. (D) Albumin was steadily removed (up to 3 mL) with a needle pointed downwards. (E) Sterile parafilm was used to cover the hole. (F) Tape placed over the parafilm to keep the parafilm secure and aid stabilisation of the egg.

*Micro-CT scan information for acquisition and reconstruction parameters*

**S. Table 2:** Settings used for µCT scanning of mice and excised samples in the subcutaneous implant studies.

| **Imaging reference** | **Samples *in vivo*** | ***Ex vivo* excised samples in sample holder** |
|---|---|---|
| Voltage (kVp) | 55 | 50 |
| Current (mA) | 0.17 | 0.21 |
| Exposure time (ms) | 75 | 65 |
| Filter (µm thickness) | Al 400+100 | Al 100 only |
| Scan angle (°) | 360 | 360 |
| Step angle (°) | 0.25 | 0.25 |
| Pixel size | 40 | 20 |
| Reconstruction voxel size (µm$^3$) | 40 | 20 |
| Frame averaging | 1 | 1 |

#### alamarBlue™ and correlation to cell number

Differing HBMSC numbers were plated in triplicate in a 24 well plate and cultured at 37 °C in 5% $CO_2$/balanced air overnight. The following day, the alamarBlue™ HS assay was performed. The results showed increased fluorescence values with increasing number of cells (**Supplementary Figure 6**).

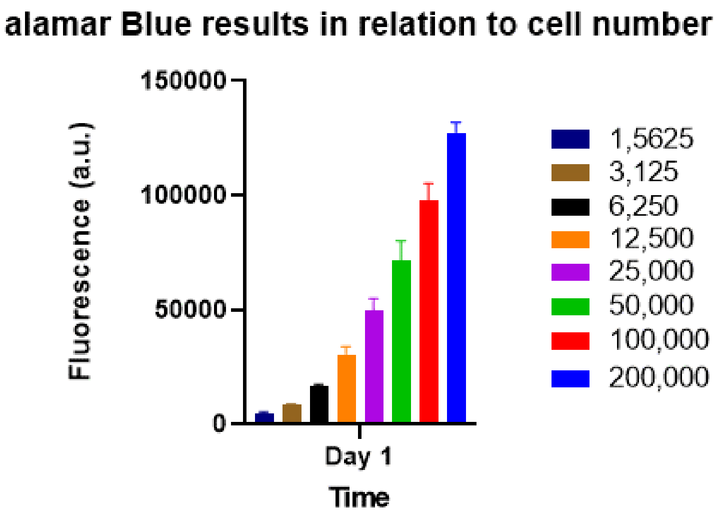


**S. Figure 6:** Differing HMBSC numbers and resultant alamarBlue™ fluorescence values. There was a trend of increasing fluorescence value as the cell number increased, approximately equating the fluorescence result to cell number. N=3, mean and S.D. shown.

#### HBMSCs were viable on PCL-TMA scaffolds with PEA/FN/BMP-2 coating

PEA/FN/BMP-2 coated PCL-TMA scaffolds were assessed compared to uncoated PCL-TMA, with uncoated nylon as the positive control. The HBMSCs adhered to the uncoated and coated PCL-TMA and multiplied over 14 days, but there was no significant difference in cell number on the scaffold at day 1 or day 14 (**Supplementary Figure 7**).

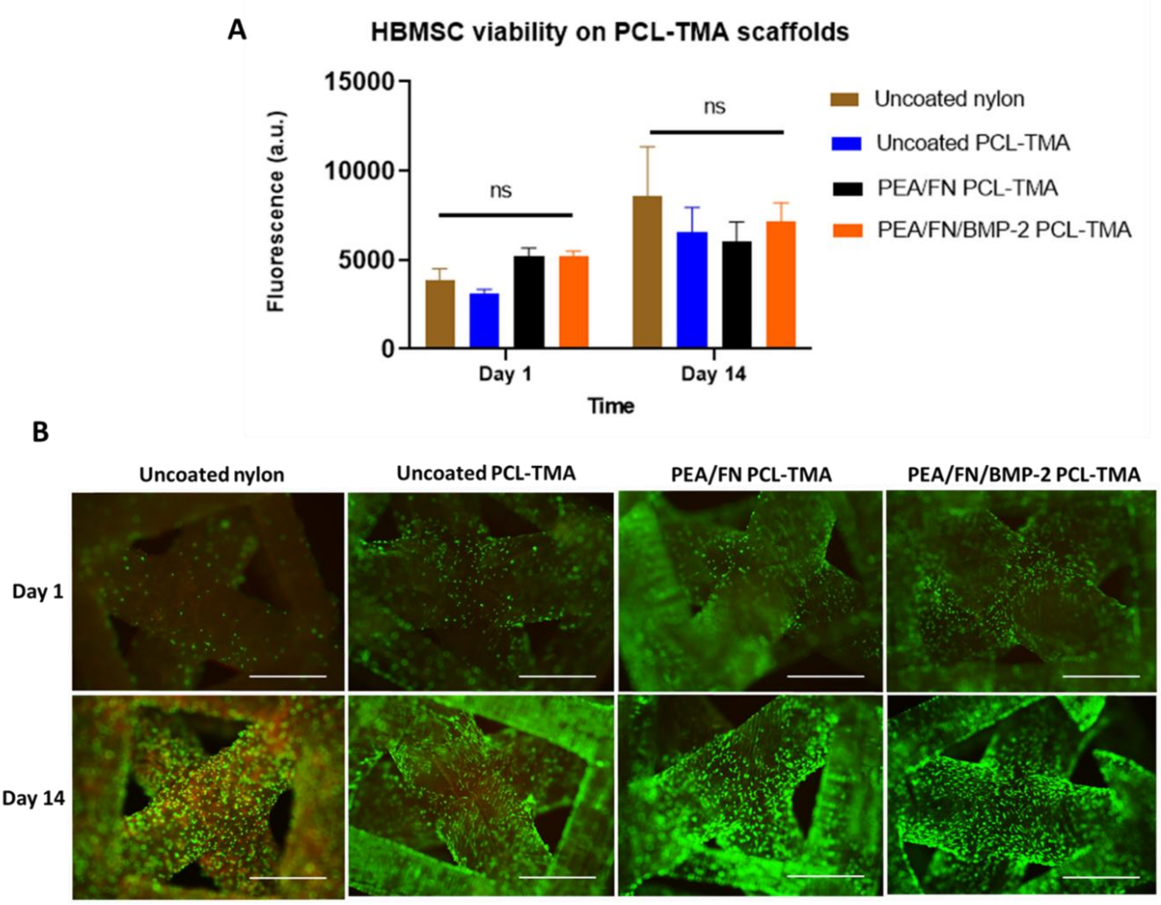


**S. Figure 7:** HBMSC viability and live (green)/dead (red) cell labelling on PCL-TMA scaffolds. (A) alamarBlue™ fluorescence results of uncoated and coated PCL-TMA scaffolds. There was no significant difference at day 1 or day 14 between PEA/FN or PEA/FN/BMP-2 coated PCL-TMA scaffolds compared to uncoated PCL-TMA. The coated scaffolds supported cell survival. (B) The cell number and location of cells adhered to the scaffold initially at day 1 and subsequent multiplication of cells over the 14-day period as seen by fluorescent labelling of live cells. Data analysed by 2-way ANOVA with Dunnett's multiple comparisons test, n=3 but n=2 for uncoated PCL-TMA and PEA/FN/BMP-2 PCL-TMA on day 1, mean and S.D. shown, ns; non-significant. Scale bar 1 mm.
